# PprA interacts with replication proteins and affects their physicochemical properties required for replication initiation in *Deinococcus radiodurans*

**DOI:** 10.1101/2020.03.25.007906

**Authors:** Ganesh K. Maurya, Hari S. Misra

## Abstract

The deletion mutant of *pprA*, a gene encoding pleiotropic functions in radioresistant bacterium *Deinococcus radiodurans*, showed an increased genomic content and ploidy in chromosome I and chromosome II. We identified *oriC* in chromosome I *(oriCI*) and demonstrated the sequence specific interaction of deinococcal DnaA (drDnaA) with *oriCI*. drDnaA and drDnaB showed ATPase activity while drDnaB catalyzed 5′→3′ dsDNA helicase activity. These proteins showed both homotypic and heterotypic interactions. The roles of C-terminal domain of drDnaA in *oriCI* binding and its stimulation of ATPase activity were demonstrated. Notably, PprA showed ~2 times higher affinity to drDnaA as compared to drDnaB and attenuated both homotypic and heterotypic interactions of these proteins. Interestingly, the ATPase activity of drDnaA but not drDnaB was inhibited in presence of PprA. These results suggested that PprA influences the physicochemical properties of drDnaA and drDnaB that are required for initiation of DNA replication at *oriCI* site in this bacterium.

## Introduction

The <u>origin of replication</u> in bacterial chromosome (*oriC*) is a discrete locus that contains AT rich conserved DNA motifs and the varying number of 9 mer repeats of non-palindromic sequences called DnaA boxes. These boxes are recognized by a replication initiator protein named DnaA, followed by the assembly of replication initiation complex at *oriC* (Messer, 2002; Mott and Berger, 2007). Mechanisms underlying replication initiation have been characterised in large number of bacteria harbouring limited copies of single circular chromosome as inheritable genetic material (Zakrzewska-Czerwińska et al., 2007; Leonard and Grimwade, 2015). It has been shown in *E. coli* that DnaA-ATP oligomer binds to DnaA boxes at *oriC* and unwinds the adjacent AT rich region. Subsequently, a hexameric complex of replicative helicase DnaB and its loader DnaC (DnaB_6_-DnaC_6_) is recruited to the unwound region in *oriC* resulting to the formation of pre-priming complex (Skarstad and Katayama, 2013; Chodavarapu and Kaguni, 2016). This provides the site for binding of primase and activation of various events required for the progression of replication complex comprised of DNA polymerase III holoenzyme. DnaB hexameric ring translocates bidirectionally in order to unwind the parental duplex DNA while 3’ end of the primer is extended by polymerase complex (McHenry, 2011; Yao and O’Donnell, 2010). In *E. coli* it has been shown that the initiation of DNA replication at *oriC* site is tightly regulated and the next round of *oriC* mediated DNA replication must wait till the *oriC* of newly replicated daughter chromosomes is fully methylated. Therefore, under normal growth conditions, the number of copies of primary chromosome per cell is expected to be less than 2 (Skarstad and Katayama, 2013). Recently, the bacteria with multiple copies of multipartite genome system have been reported. Notably, most of them are either parasite to some forms of life or exhibits super tolerance to abiotic stresses (Misra et al, 2018). The ploidy of chromosomes in these bacteria allowed us to revisit the principle of *oriC* regulation as known in bacteria containing less than 2 copies of circular chromosome per cell. Mechanisms underlying regulation of *oriC* function in multipartite genome harbouring bacteria have not been studied in detail. In case of *Vibrio cholera*, which harbours two chromosomes viz; chromosome I (Chr I) and chromosome II (Chr II), the Chr I replicates similar to *E. coli* chromosome while Chr II follows replication mechanism similar to low copy number P1 and F plasmids (Egan et al., 2005; Egan and Waldor, 2003; Jha et al., 2012; Ramachandran et al., 2017; Koch et al., 2010; Fournes et al., 2018).

*Deinococcus radiodurans* is one of the most radiation resistant bacteria, characterized for its extraordinary ability to withstand the lethal effects of DNA damaging agents including radiation and desiccation (Zahradka et al., 2006; Cox and Battista, 2005; Slade and Radman, 2011; Misra et al., 2013). It also harbours the multipartite genome system comprised of two chromosomes (Chr I (2,648,638bp) and ChrII (412,348bp) and a megaplasmid (177,466bp) and plasmid (45,704 bp) (White et al., 1999). Interestingly, each of these genome elements is found in multiple copies per cell (Hansen, 1978). Chr I of *D. radiodurans* encodes putative DnaA (DR_0002) and DnaB (DR_0549) (hereafter named drDnaA and drDnaB, respectively) while chromosome II encodes PprA (DR_A0346), which has been characterized for various functions (Narumi et al., 2004, Kota et al., 2014, 2016; Adachi et al, 2014). Recently, the extended structure of PprA has been reported where a possibility of it acting as protein scaffold (Adachi et al., 2019), which might explain PprA roles in large number of molecular events. Here, for the first time, we report the functional characterization of chromosome replication initiation proteins drDnaA and drDnaB in *D. radiodurans* and demonstrated that PprA a pleiotropic protein involved in radioresistance plays a role in the regulation of DNA replication. We showed that drDnaA interacts with *origin of replication* (*oriCI*) in a sequence specific manner. drDnaA is characterized as an *oriCI* responsive ATPase while drDnaB as ATP dependent 5′→3′ dsDNA helicase with higher affinity for ssDNA as compared to dsDNA. Interestingly, PprA interacted with drDnaA at relatively higher affinity than drDnaB and inhibited both homotypic and heterotypic interactions of these proteins. Further, PprA downregulated the ATPase activity of drDnaA but showed no effect on ATPase and helicase activities of drDnaB. These results suggested that drDnaA and drDnaB confers the desired functions for the initiation of replication at *oriCI* in *D. radiodurans.* Furthermore, the interference of PprA in the physicochemical properties of these replication proteins and an increase in copy numbers of both primary and secondary chromosomes in absence of *pprA* has suggested its involvement in the regulation of chromosomal replication in this bacterium.

## Results

### The *pprA* deletion affects genomic content in *D. radiodurans*

Earlier, the regulatory role of PprA in cell division and genome maintenance has been demonstrated (Devigne et al., 2013; Kota et al., 2014; Devigne at al., 2016; Kota et al., 2016). The DNA content and the copy number of genome elements in *pprA* deletion mutant were compared with wild type *D. radiodurans*. The total DNA content in mutant cells (~8.41 ± 1.05 fg per cell) was found to be ~3 fold higher than wild type cells (~3.05 ± 0.47 fg per cell) (Fig. 1A). Similarly, the DAPI fluorescence in Δ*pprA* mutant was ~2 fold higher as compared to wild type (Fig. 1B). The cell scan analysis of DAPI stained cells showed that ~80 % Δ*pprA* cells have ~2 folds higher DAPI fluorescence as compared to wild type cells grown identically (Fig. 1C). When we checked the copy number of each replicon, the average copy number of Chr I and Chr II was ~2.5 to 3-fold higher in Δ*pprA* mutant as compared to wild type. For instance, the copy number of Chr I was ~ 8 in wild type while it was ~18 in Δ*pprA* mutant. The copy number of Chr II had also increased from ~7 in wild type to ~17 per cell in Δ*pprA* mutant (Fig. 1D). Interestingly, the copy number of megaplasmid and small plasmid did not change. This increase in genomic content and chromosomal copy number indicated a strong possibility of continued DNA replication in the absence of PprA. Thus, a possible role of PprA in the regulation of DNA replication is suggested in *D. radiodurans*.

**Fig. 1.**
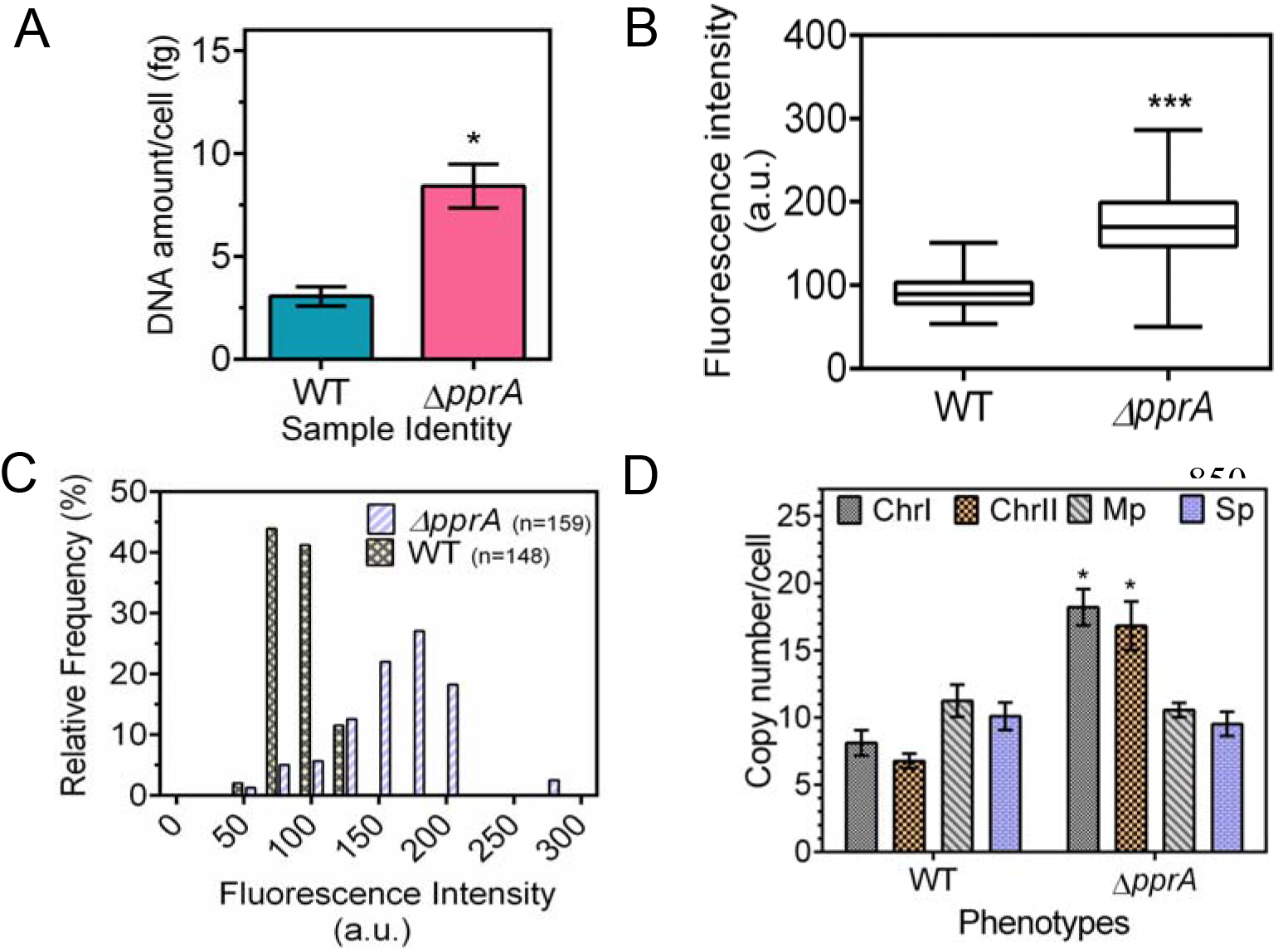
Change in genomic content in *pprA* deletion mutant of *D. radiodurans*. *D. radiodurans* R1 (WT) and its *pprA* mutant (Δ*pprA*) were grown to mid-logarithmic phase. The amount of DNA per cell was measured spectrophotometrically (A) and through DAPI staining of nucleoid (B). The change in DAPI fluorescence in wild type and Δ*pprA* mutant cells was scanned in ~150 cells and relative frequency of cells containing different amount of DAPI stained nucleoid was estimated (C). Similarly, wild type and mutant populations were determined for the copy number of chromosome I (ChrI), chromosome II (ChrII), megaplasmid (Mp) and small plasmid (Sp) (D). Data presented in A, B and D are mean ± SD (n=9). Data given in C are from the wild type cell population (n=148) and mutant population (n=159).

### PprA interacts with DnaA and DnaB of *D. radiodurans*

In order to get the mechanistic insights on the regulatory role of PprA in DNA replication, the physical and functional interaction of PprA with drDnaA and drDnaB proteins were separately monitored. *E. coli* BTH101 (*cyaA*^−^) co-expressing T18 tagged PprA and T25 tagged drDnaA or drDnaB showed reconstitution of active CyaA leading to the expression of β-galactosidase activity (Fig. 2A). The reconstitution of active CyaA from its T18 and T25 domains would be possible only when target proteins tagged with these domains interact. PprA interaction with drDnaA and drDnaB was further confirmed using surface plasmon resonance. For that recombinant PprA protein was immobilized on a bare gold sensor chip and the purified recombinant drDnaA or drDnaB (Fig. S2) was passed over the immobilised PprA in the mobile phase. Results showed a concentration-dependent increase in SPR signals in case of drDnaA or drDnaB (Fig. 2 B,C). The dissociation constant (Kd) for drDnaA was 5.41 × 10^−7^ ± 1.8 × 10^−8^ [M] while it was 9.71 × 10^−7^ ± 1.1 × 10^−8^ [M] for drDnaB. These indicated that PprA interacts with drDnaA with ~2-fold higher affinity than drDnaB. We have also tested the interaction of C-terminal deletion mutant of DnaA (DnaΔCt) with PprA in surrogate *E. coli* using co-immunoprecipitation, as described in methodology. We found that truncation of C-terminus from drDnaA has not affected its interaction with PprA (Fig. 2D). This suggests a possibility of N-terminal and/or middle region of drDnaA could be the region of interaction with PprA protein in this bacterium.

**Fig 2.**
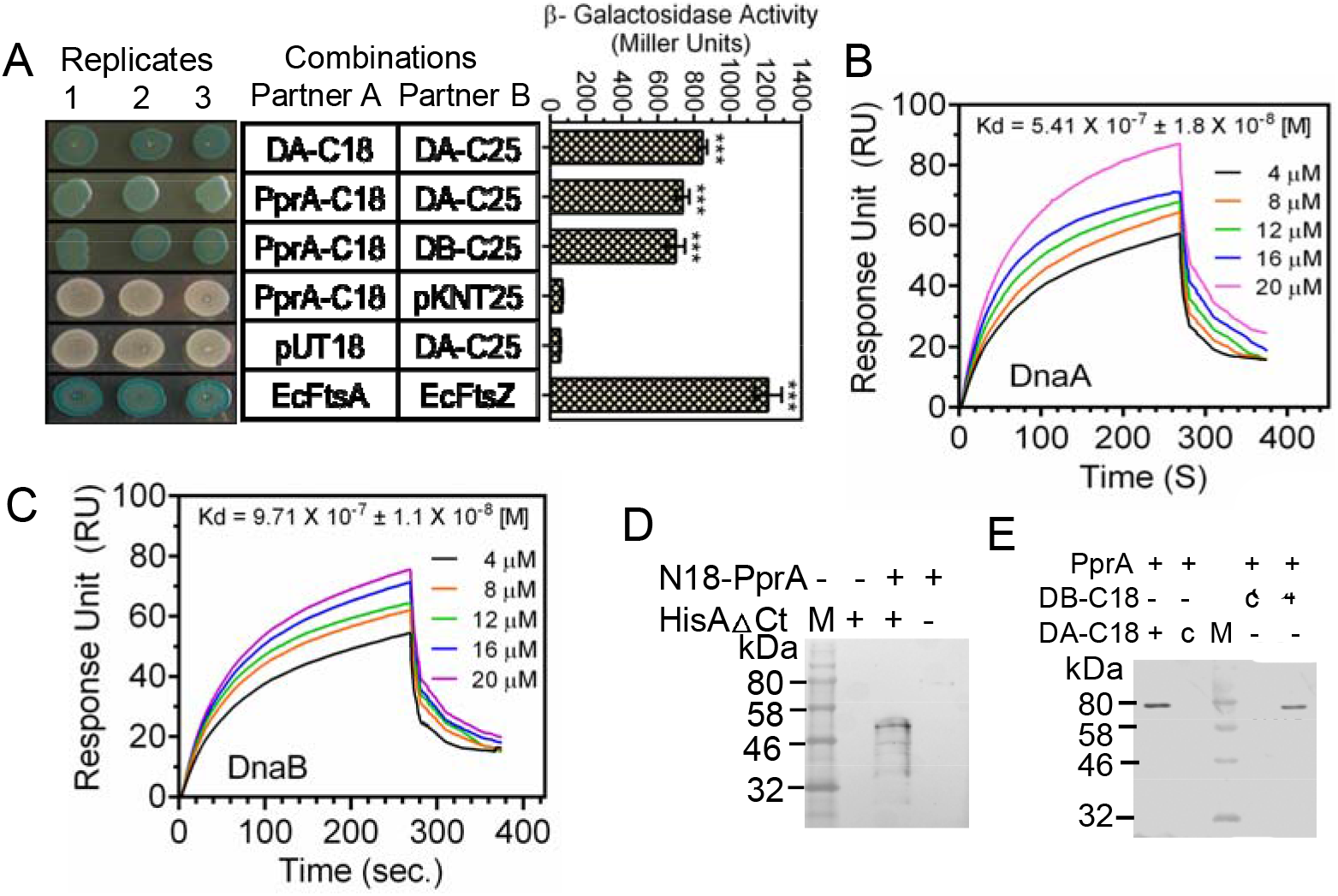
PprA interaction with drDnaA and drDnaB of *D. radiodurans*. *E. coli* BTH 101 cells co-expressing drDnaA-C18 (DA-C18), drDnaA-C25 (DA-C25), drDnaB-C25 (DB-C25), PprA-C18 in different combinations were detected for expression of β-galactosidase in spot test and in liquid (A). The bacterial two hybrid system vectors pUT18 and pKNT25 were used as negative control while *E. coli* FtsA (EcFtsA) and FtsZ (EcFtsZ) were used a positive control. Recombinant PprA interactions with recombinant drDnaA (B) and drDnaB (C) of *D. radiodurans* were studied by surface plasmon resonance. *Ex vivo* interaction of PprA expressing from PprA-C18 and C-terminal truncated DnaA with histidine tag (HisAΔCt) was monitored in surrogate *E. coli* (D). *In vivo* interaction of native PprA expressing on chromosome and drDnaA (DA-C18), and drDnaB (DB-C18) on plasmids was monitored in *D. radiodurans* R1. Total proteins were precipitated with antibodies against polyhistidine (D) and PprA (E). Perspective interacting partners were detected using antibodies against T18 domain of CyaA. Sizes of fusions were compared with molecular weight marker (M). C in panel E indicates plasmid control.

The interaction of PprA with drDnaA or drDnaB was also monitored in *D. radiodurans*. For that, the total proteins of *D. radiodurans* cells expressing T18 tagged drDnaA or drDnaB on plasmid and PprA on chromosome were precipitated using antibodies against PprA, and interaction of PprA with drDnaA / drDnaB partners if any, were detected using T18 antibodies. PprA interaction with drDnaA and drDnaB was confirmed in *D. radiodurans* (Fig. 2E). These results suggested that PprA interacts with both drDnaA and drDnaB in *D. radiodurans*, and that might help PprA to regulate the functions of these proteins in this bacterium.

### The chromosome I of *D. radiodurans* contains putative *oriCI*

The putative origin of replication of chromosome I in *D. radiodurans* R1 was predicted using DOriC database (DoriC accession number - ORI10010007) (Luo and Gao, 2019). The 1183 to 1903 bp upstream to dr*dnaA* gene (DR_0002) was searched for consensus sequences of 9 mer DnaA-boxes using WebLogo online tool (Crooks et al., 2004). The 1273 to 1772 bp (~500 bp) region upstream to dr*dnaA* coding sequence contains 13 DnaA-boxes of 9 mer each and was named as putative *oriCI*. These DnaA boxes are distributed in random fashion on both sense (4 DnaA boxes) and antisense (9 DnaA boxes) strands of chromosome I (Fig. 3). Unlike *E. coli oriC*, the structure of *oriCI* in *D. radiodurans* is complex. For instance, it is comparatively longer than the *oriC* of *E. coli* and contains many ‘non-perfect’ DnaA-boxes. The *oriCI* has 46.2 % GC content indicating that it is an AT rich region. When we analysed this region for typical *E. coli* type DnaA-boxes, 8 out of 13 DnaA-boxes were perfect like DnaA-boxes (TTATCCACA) of *oriC* in *E. coli* (Messer, 2002). The other 5 boxes differ from *E. coli* type by single nucleotide (Fig. 3) but were like the DnaA-boxes in the chromosome of other bacteria like *Cyanothece* 51142, *T. thermophilus* and *B. subtilis* (Huang et al., 2015; Schaper et al., 2000; Moriya et al., 1988).

**Fig. 3.**
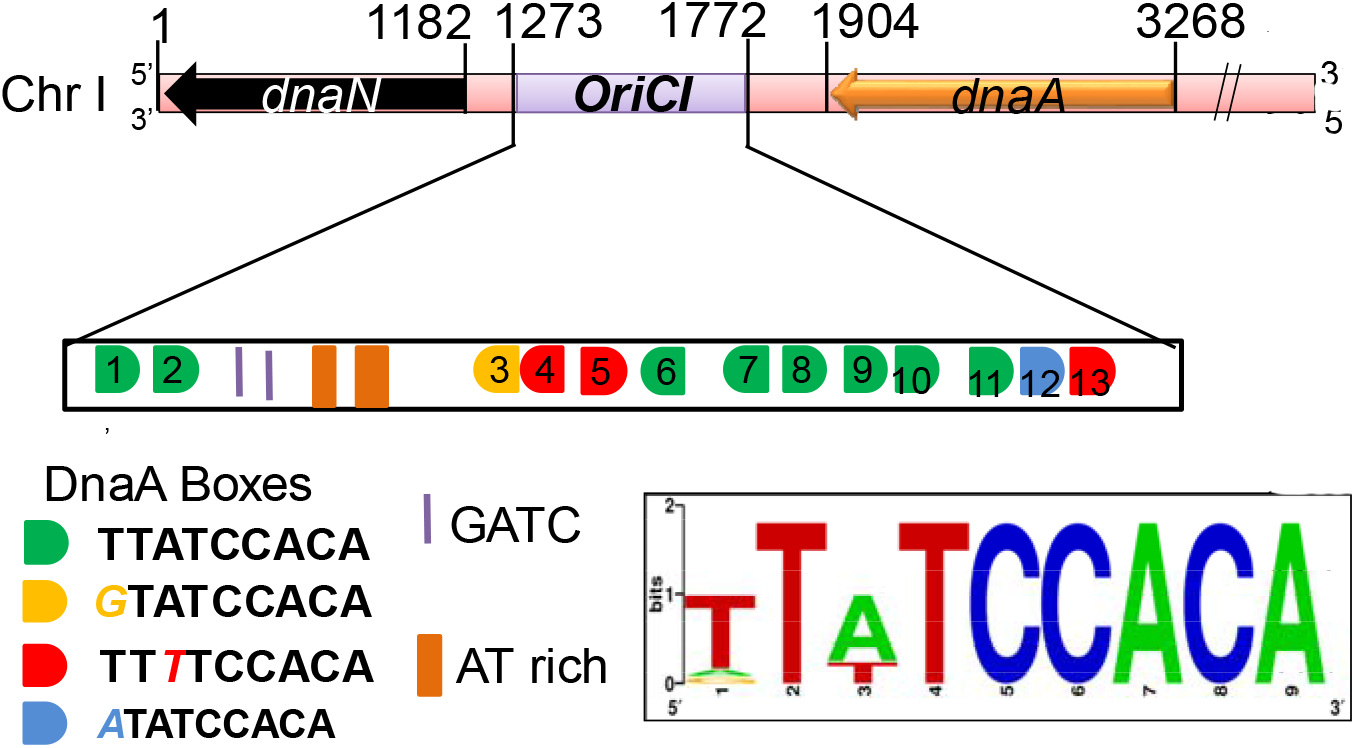
Organization of putative origin of replication in chromosome I (*oriCI*) of *D. radiodurans.* A *cis* element region (1273-1772) located between *dnaN* and *dnaA* in chromosome I of *D. radiodurans* was analyzed for putative *oriC* signature. This fragment is found containing 13 copies of conserved DnaA boxes, 2 GATC motif and 2 AT rich regions and thus predicted to be *oriCI in D. radiodurans*.

A web logo of these DnaA-boxes has generated a consensus sequence of T(A/G)TA(T)TCCACA. These DnaA-boxes are distributed in random fashion on both sense (4 DnaA-boxes) and antisense (9 DnaA-boxes) strands of chromosome I (Fig. 3). This suggested that *oriCI* in chromosome I of *D. radiodurans* is largely similar to *oriC* of *E.coli*.

### drDnaA is a sequence specific *oriCI* binding protein

The recombinant drDnaA was purified from recombinant *E. coli* (Fig.S1A) and its *oriCI* binding activity was checked by electrophoretic mobility shift assays (EMSA). *oriCI* containing 13 repeats of DnaA-boxes was generated by PCR amplification and radiolabelled using [^32^P]γ-ATP. The DNA substrate was incubated with purified recombinant drDnaA and the nucleoprotein complex was separated on native poly-acrylamide gel electrophoresis (PAGE). drDnaA showed sequence-specific interaction with *oriCI in vitro* as this interaction was not affected by addition of 50-fold excess of cold non-specific DNA (Fig. 4A, B). Interestingly, the affinity of drDnaA to *oriCI* (Kd = 1.68 ± 0.23 μM) increased by ~ 8-fold in the presence of ATP (Kd = 0.243 ± 0.02 μM) (Fig. 4C, D). We further checked the affinity of DnaA with different number of DnaA boxes in *oriCI* in the presence of ATP and found that as the number of DnaA boxes reduces, the affinity of DrDnaA also reduces (Table 1). . Like *E. coli* DnaA, drDnaA has also shown specific affinity for single DnaA box though the affinity with perfect DnaA box (TTATCCACA; Kd = 2.88 ± 0.28 μM) is ~2 fold higher than non-perfect type DnaA boxes like TTTTCCACA or GTATCCACA (Kd = 4.21 ± 0.15 μM or 4.36 ± 0.18 μM, respectively) (Fig. 2 G-I).These results showed that drDnaA is an *oriCI* binding protein and its interaction with *oriCI* is stimulated in the presence of ATP.

**Fig. 4.**
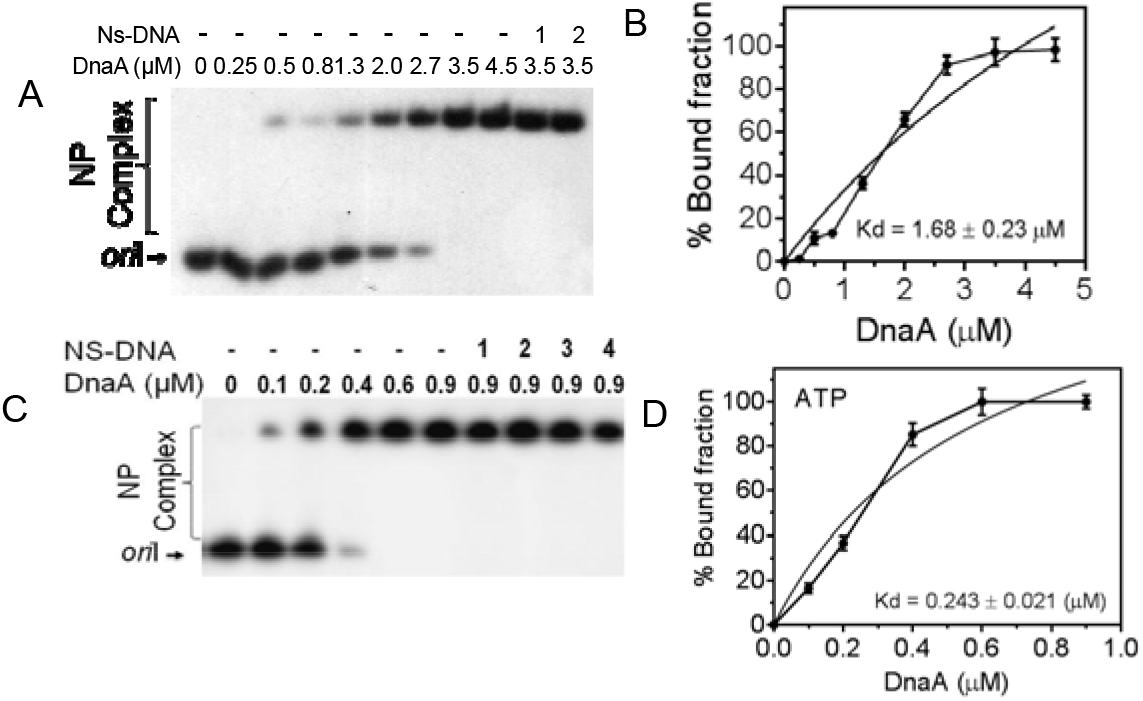
Recombinant drDnaA interaction with putative *oriCI*. The *oriCI* (*oriI*) DNA was radiolabeled and incubated with increasing concentration of recombinant drDnaA protein in the absence (A, B) and presence (C, D) of 1mM ATP. The nucleoprotein complexes were analyzed on native PAGE. The 5 (1), 10 (2), 20 (3) and 50 (4) fold higher molar concentration of non-specific dsDNA (NS-DNA) was added into reaction mixture and analyzed on non-denaturing PAGE. Band intensity of free form and bound form of DNA was estimated densitometrically and plotted as mean ± SD (n=3) of bound fraction in percentage.

**Table 1.** Dissociation constant (Kd) of drDnaA with *oriI* and its repeat variants with different number of DnaA boxes was measured in the presence and absence of ATP.

| Number of DnaA boxes | ATP | Dissociation constant (Kd) (μM) |
| --- | --- | --- |
| 13 (TTATCCACA)<sub>13</sub> | - | 1.78±0.23 |
| 13 (TTATCCACA)<sub>13</sub> | + | 0.243±0.021 |
| 11(TTATCCACA)<sub>11</sub> | + | 0.241±0.01 |
| 7(TTATCCACA)<sub>7</sub> | + | 0.373±0.01 |
| 3(TTATCCACA)<sub>3</sub> | + | 1.08±0.12 |
| 1 (TTATCCACA) | + | 2.88±0.28 |
| 1 (TTTTCCACA) non-perfect repeat | + | 4.21±0.15 |
| 1 (GTATCCACA) non-perfect repeat | + | 4.36±0.18 |

### drDnaA binds *oriCI* through its C-terminal and N-terminal domain associated ATPase activity is stimulated by *oriCI in vitro*

Since drDnaA showed relatively higher affinity for *oriCI* in the presence of ATP (Fig. 4C), the ability of drDnaA for ATP hydrolysis and the effect of *oriCI* on its ATPase function was further investigated. We observed that drDnaA can hydrolyse [^32^P] αATP into [^32^P] αADP (Fig 5A) and this activity was stimulated in the presence of *oriCI* (Fig. 5B, 5C). Interestingly, the ATPase activity of drDnaA was not affected in the presence of non-specific dsDNA (Fig. S3). The helix-turn-helix motif in domain IV at the C-terminal of DnaA has been reported for sequence specificity in many bacteria (Roth and Messer, 1995; Sutton and Kaguni, 1997; Majka et al., 1999; Blaesing et al., 2000; Messer et al., 1999; Fujikawa et al., 2003). Therefore, the C-terminal (domain IV) truncation was created in drDnaA (DnaAΔCt) (Fig 6A) and its binding to *oriCI* was checked in the presence of ATP. DnaAΔCt failed to bind *oriCI* (Fig. 6B). Although, this truncation did not affect ATPase activity of drDnaA (Fig 6C), as expected the *oriCI* stimulation of ATPase activity was not observed in DnaAΔCt protein (Fig 6D, 6E). These results suggested that drDnaA is an *oriCI* binding ATPase and its C-terminal domain IV is essential for recognition to *oriCI* sequence and thereby the ATPase activity stimulation by this cognate *cis* element.

**Fig. 5.**
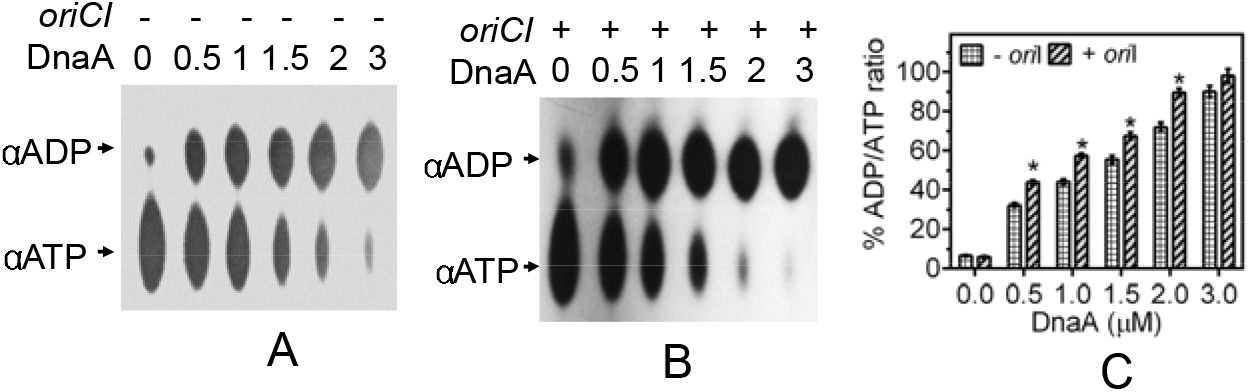
ATPase activity in recombinant drDnaA. An increasing concentration (0.5, 1, 1.5, 2.0 and 3 μM) of recombinant drDnaA (DnaA) was incubated with [^32^P]-αATP (αATP) in the absence (A) and presence (B) of *oriCI* (*oriI*)and generation of [^32^P]-αADP (αADP) product was detected on TLC after autoradiography. Spot intensity was quantified densitometrically and percent of ADP/ATP ratios were plotted as a function of drDnaA concentration (C). Results were analyzed using student t-test and significant difference in data sets with p values of 0.05 or less is shown as (*).

**Fig. 6.**
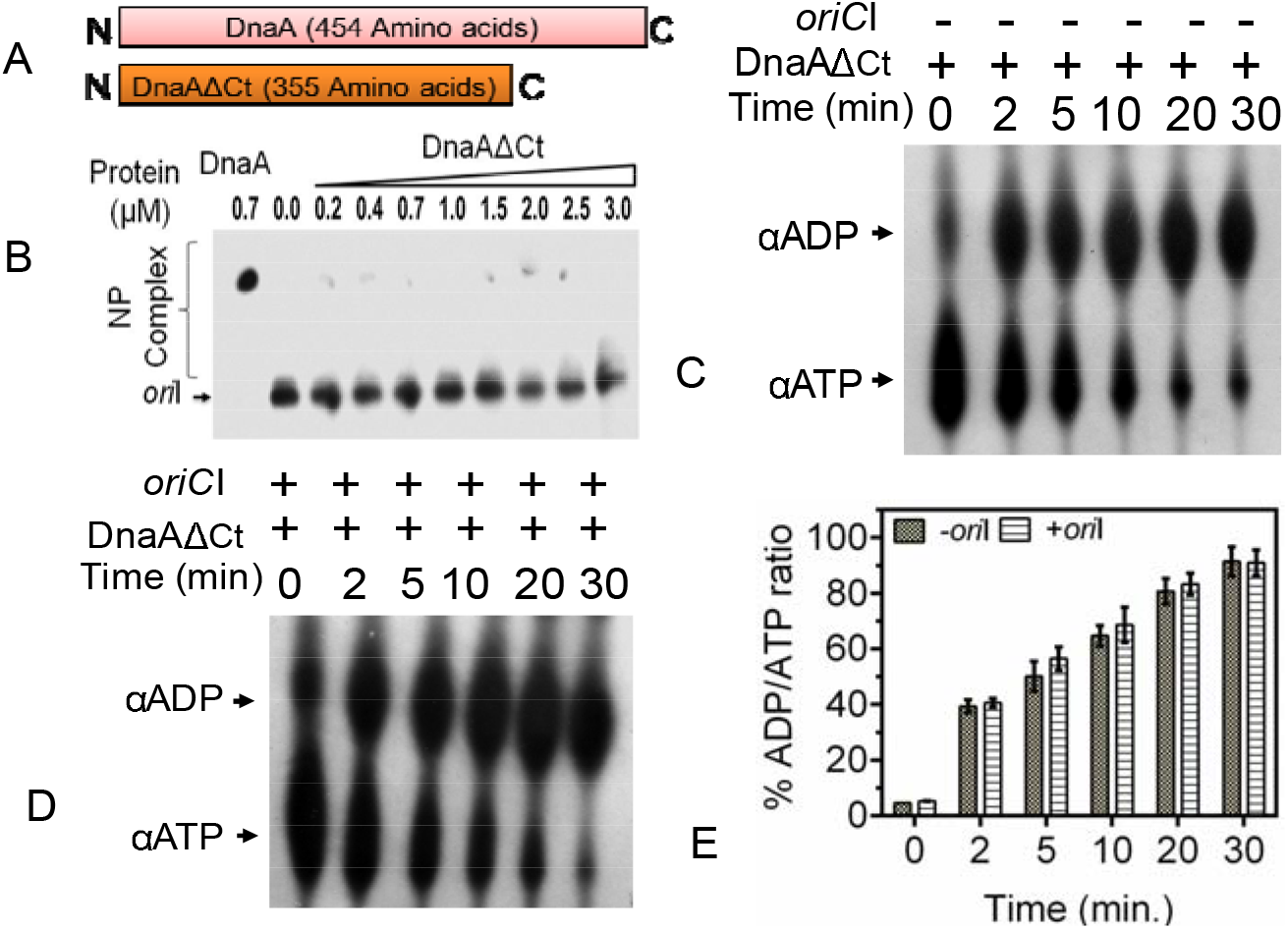
Role of C-terminal putative helix turn helix motifs containing domain in drDnaA function. The 99 amino acids of drDnaA were removed from its C-terminal and resulting DnaAΔCt derivative was generated (A). An increasing concentration of recombinant protein was incubated with radiolabeled *oriCI* (*oriI*) as described in Fig 4A. Products were analysed on native PAGE and autoradiogram was developed (B). Similarly, 2 μM concentration of DnaAΔCt was incubated for different time points with [^32^P]-αATP in the absence (C) and presence (D) of *oriCI* (*oriI*). Products were separated on TLC and analyzed as described in Fig 5. The percentage of ADP/ATP ratios were plotted as a function of time (E). Results were analyzed using student t-test and significant difference in data sets with p values of 0.05 or less is shown as (*).

### drDnaB is a ssDNA responsive ATPase and an ATP dependent 5′→3′ dsDNA helicase

drDnaB showed non-specific DNA binding activity with *oriCI* as chase with non-specific dsDNA has significantly competed out *oriCI* from drDnaB binding. The affinity of drDnaB to *oriCI* (Kd = 8.89±0.92 μM) was not affected by ATP (Kd = 8.23±0.41 μM). However, drDnaB affinity to ssDNA (Kd= 3.11±0.11 μM) was nearly ~3 times higher than dsDNA (*oriCI*) (Kd= 8.89±0.92 μM). drDnaB affinity to ssDNA was further increased by 2 folds in the presence of ATP (Kd = 1.42±0.05 μM) (Fig. 7). These results suggested that drDnaB prefers binding to ssDNA over dsDNA in ATP independent manner, and addition of ATP improves its affinity for ssDNA but not to dsDNA. drDnaB could hydrolyse [^32^P]-αATP to [^32^P]-αADP (Fig 8A), which was stimulated in the presence of ssDNA (Fig. 8B, 8C). Similar observations were reported earlier for other DnaB homologs (Biswas et al., 2009; Zhang et al., 2014). DnaBs are characterized as the replicative helicases and play essential role in *oriC* mediated DNA replication in other bacteria (LeBowitz and McMacke, 1986; Patel and Picha, 2000; Zawilak, et al., 2001). Therefore, the helicase activity of recombinant drDnaB was checked on dsDNA having either 3′ or 5′ overhangs in the presence and absence of ATP and products were monitored in gel and FRET based assays. Results showed that drDnaB unwinds dsDNA with 5′ overhang only and not with 3′ overhang (Compare Fig. 9A and 9E with 9C and 9F). This 5′ → 3′ dsDNA helicase activity required the hydrolysable ATP as no DNA unwinding was observed when ATP was replaced with ATP-γ-S (Fig 9B). The similar results were obtained in FRET assays with FAM as donor and BHQ as acceptor (Fig 9D). Results showed that the fluorescence has increased in the presence of protein and ATP but not with ATP-γ-S (Fig 9E and 9F). These results suggested that drDnaB is an ATP dependent 5′ → 3′ dsDNA helicase, which largely agrees with the characteristics of DnaBs of other bacteria (Soultanas and Wigley, 2002; Soni et al., 2003; Nitharwal et al., 2012; Zhang et al., 2014).

**Fig. 7.**
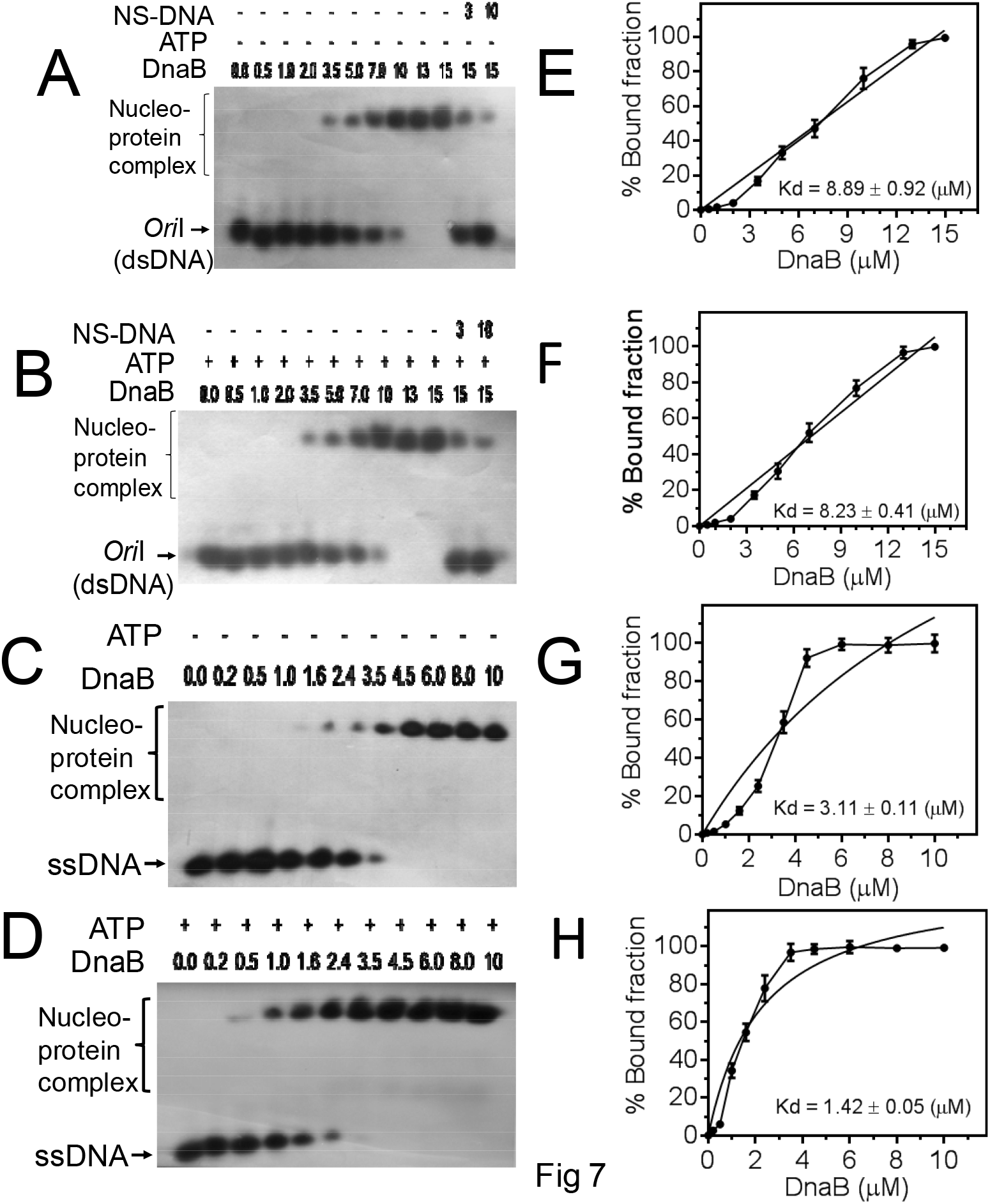
DNA binding activity of DnaB of *D. radiodurans* (drDnaB). An increasing concentration (μM) of recombinant purified drDnaB (DnaB) was incubated with radiolebeled *oriCI* (*oriI*) taken as dsDNA (A, B) and radiolabeled ssDNA (C and D) in the absence (A and C) and presence (B and D) of ATP. Products were analyzed on non-denaturing PAGE. Autoradiograms were developed, band intensity of free and bound form of DNA was estimated densitometrically from autoradiogram A (E), B (F), C (G) and D (H), and the mean ± SD (n=3) of percent bound fraction was plotted.

**Fig. 8.**
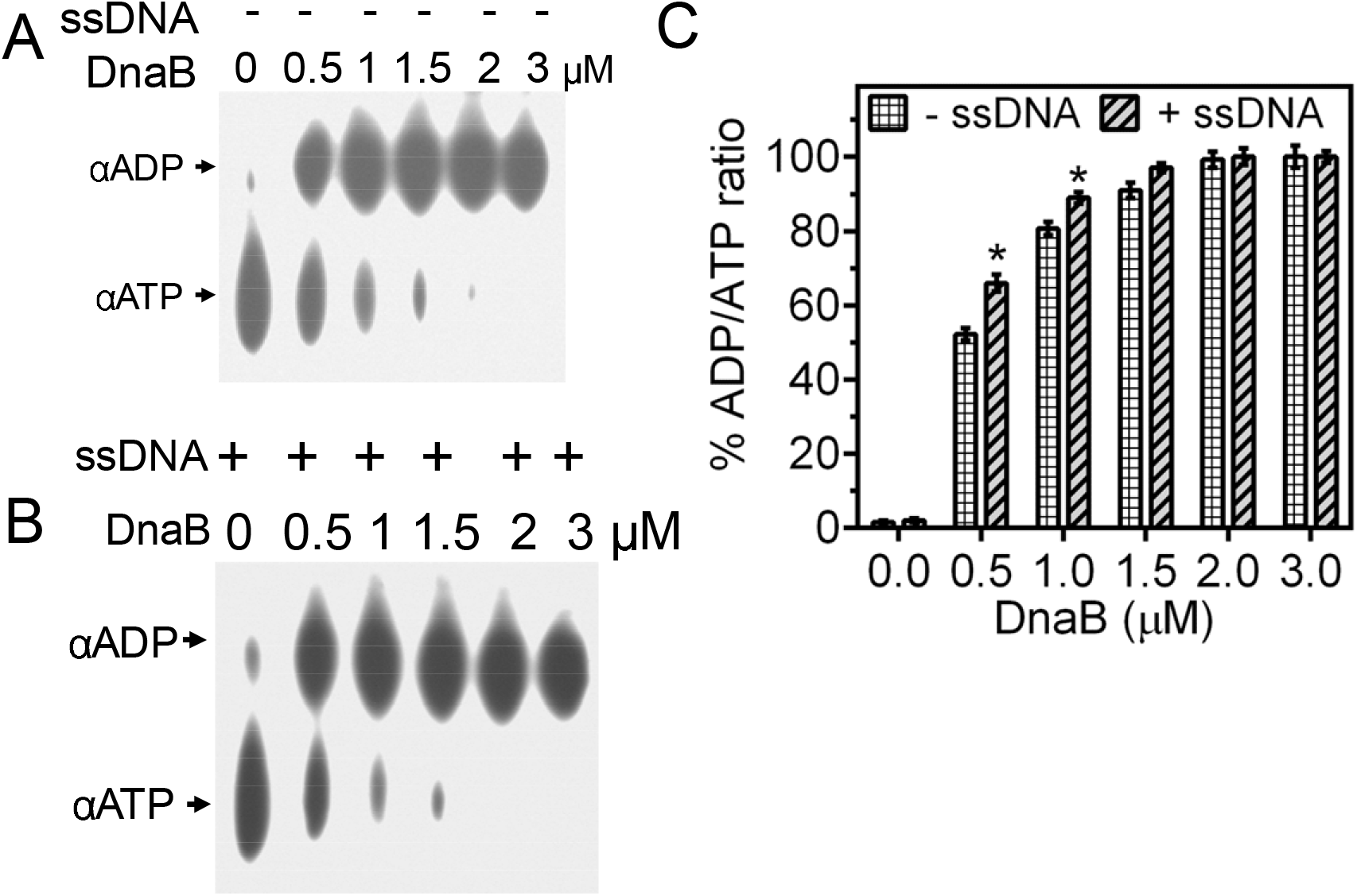
ATPase activity of purified recombinant drDnaB. An increasing concentration of drDnaB (DnaB) was incubated with [^32^P]-αATP (αATP) in the absence (A) and presence (B) of ssDNA and generation of [^32^P]-αADP (αADP) product was detected on TLC. Spot intensity was quantified densitometrically and the percent of ADP/ATP ratios were plotted as mean ± SD (n=3). Results were analyzed using student t test and significant difference in data set with p values of 0.05 or less is marked with (*).

**Fig. 9.**
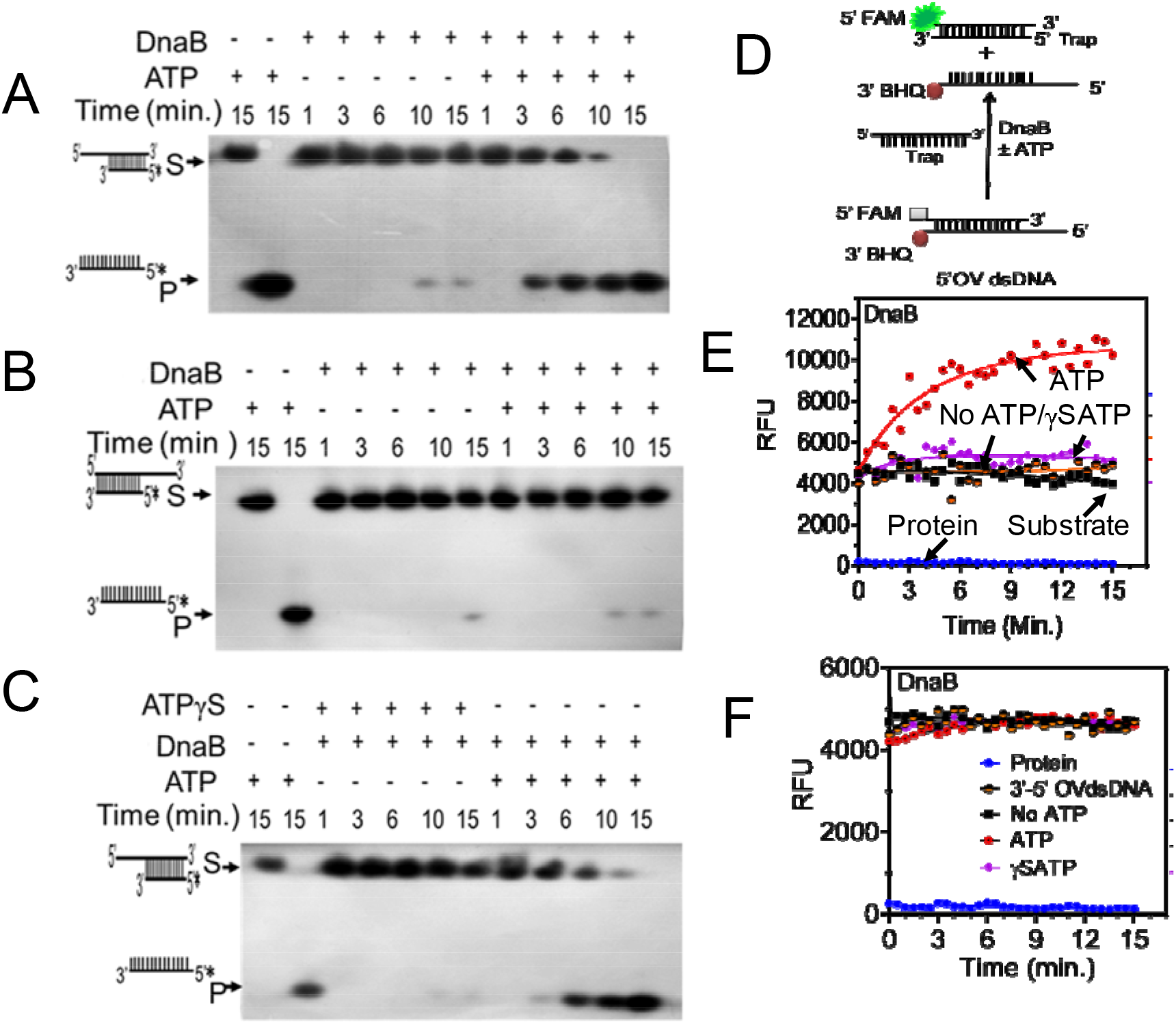
Helicase activity assay of recombinant drDnaB. The purified recombinant drDnaB (DnaB) (2 μM) was incubated with radiolabeled dsDNA having 5’ overhang (A) and 3’ overhang (B) in the presence and absence of ATP. The role of ATP hydrolysis on helicase activity was monitored with dsDNA with 5’ overhang in the presence and absence of ATP and ATP-γ-S for different time points (C). Products were analyzed on denaturing PAGE and autoradiograms were developed. The helicase activity was also monitored using FRET assay on substrate containing FAM at 5’ end and BHQ at 3’ end of dsDNA (D). Loss of FRET was monitored as gain of FAM emission with the substrate having 5’ overhang (E) and 3’ overhang (F) as a function of time in the presence and absence of ATP and ATP-γ-S. Data represents the sets of the reproducible experiment repeated 3 times.

### drDnaA and drDnaB show homotypic and heterotypic interactions

The possible interactions of drDnaA and drDnaB proteins were monitored using bacterial two-hybrid (BACTH) system and co-immunoprecipitation *ex-vivo* and *in vivo*. Both N-terminal and C-terminal fusions of drDnaA and drDnaB with T18 and T25 BACTH tags were expressed in *E. coli* BTH101 (*cyaA*^−^) (Fig S4), and the expression of β-galactosidase upon reconstitution of active CyaA from its T18 and T25 domains through interactions of target proteins was monitored. The *E. coli* expressing drDnaA and drDnaB tagged with T18 and T25 through their C-terminals showed both homotypic and heterotypic interactions and the expression of β-galactosidase activity was observed in both spot and liquid assay (Fig. 10). Some of these interactions were further confirmed by immunoprecipitation in *E. coli* BTH101 as described in method. The co-IP results from surrogate *E. coli* agreed with the results obtained from BACTH analysis (Fig 10 B,C). For *in vivo* interaction studies, the *D. radiodurans* cells were co-expressed drDnaA or drDnaB with T18 tag and / or polyhistidine tag on respective plasmids. Total proteins from these cells were immunoprecipitated using polyhis antibodies and the probable interacting partner was detected using anti-T18 antibodies. The results obtained were similar to that of BACTH and co-IP analysis in surrogate *E. coli* BTH101 expressing these proteins in different combination (Fig. 10 D). The results indicated that both these proteins interact through N-terminal domains as reported for their homologs (Seitz et al., 2000; Messer, 2002; Matthew and Simmons, 2019). Since, C-terminal domain of drDnaA is required for *oriCI* binding but not for both homotypic and heterotypic interaction with drDnaA and drDnaB, the interaction of these proteins seems to be independent of drDnaA interaction to *oriCI*. This suggested that the N-terminal and or middle region of these proteins seems to involve in oligomerization. This was further confirmed when DnaAΔCt did not affect its interaction with full length drDnaA and drDnaB (Fig. 10 E). Further mutational analysis of both deinococcal DnaA and DnaB would require to exactly map the interaction domains or residues between these proteins.

**Fig 10.**
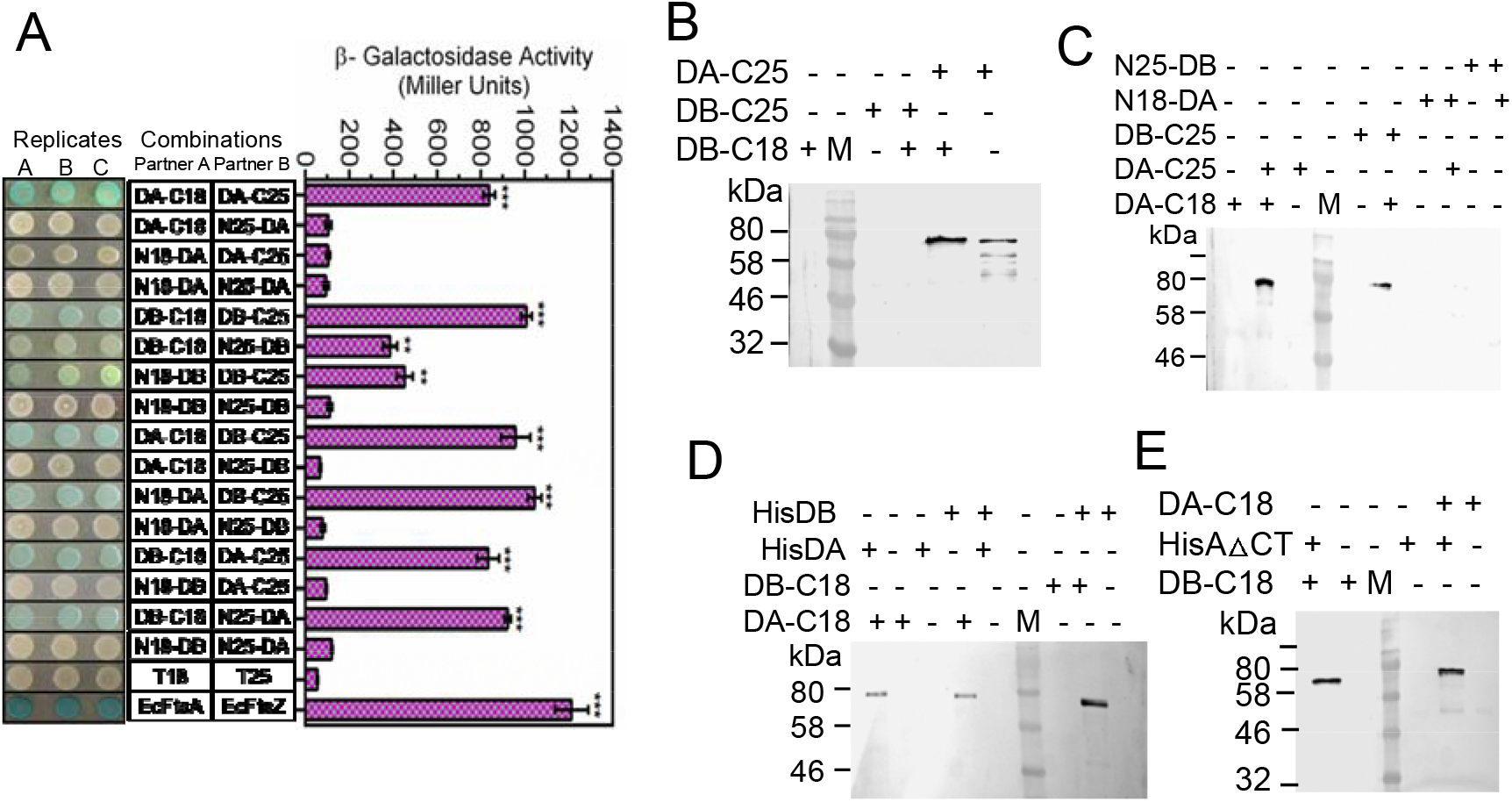
Interaction between drDnaA and drDnaB proteins. The translation fusions of drDnaA and drDnaB with T18/T25 tags at N-terminal (N18/N25-DA and N18/N25-DB) and at C terminal (DA-C18/C25 and DB-C18/C25) were expressed in *E. coli* BTH 101 cells in different combination. The expression of β-galactosidase was checked in spot test and in liquid culture (A). Total proteins from *E. coli* expressing some of these combinations were immunoprecipitated with T25 antibodies and interacting partner were detected using T18 antibodies (B and C). The pUT18 (T18) and pKNT25 (T25) vector were used as negative control while *E. coli* FtsA (EcFtsA) and FtsZ (EcFtsZ) were used a positive control. *In vivo* interaction of drDnaA (DA-C18), DnaA with histidine tag (HisDA), drDnaB (DB-C18), drDnaB with histidine tag (HisDB) and C-terminal truncated drDnaA with histidine tag (HisDAΔct) was monitored in *D. radiodurans* R1 (WT) expressing these recombinant proteins in different combinations on the plasmids. Total proteins were precipitated with polyhis antibodies and perspective interacting partners were detected using antibodies against T18 domain of CyaA.

### PprA modulates *in vitro* function of drDnaA but not drDnaB

The ATP hydrolysis is the key feature of bacterial DnaA and DnaB proteins characterized till date, and this activity is required for the initiation and progression of *oriC* mediated replication. drDnaA and drDnaB proteins exhibit ATPase activity *in vitro* (Fig. 5, 8). Therefore, the effect of PprA *per se* on ATPase activity of cognate drDnaA and drDnaB proteins was checked *in vitro*. Interestingly, PprA did not hydrolyse ATP into ADP by itself. However, the ATPase activity of drDnaA was significantly reduced in the presence of equimolar concentration of PprA irrespective of the order of addition of these proteins in reaction mixture (Fig. 11 A, B).There was no effect of PprA on either ATPase activity (Fig 11C & 11D) or 5′→3′ dsDNA helicase activity (Fig 12) of drDnaB *in vitro*. Since, PprA did not affect drDnaB function *in vitro*, the possibility of ATP titration by PprA and if that becomes the limiting factor for ATPase activity of drDnaA was nearly ruled out. Since the ATPase activity of DnaA is required for *oriC* dependent replication initiation in bacteria (Messer, 2002; Chodavarapu & Kaguni, 2016), the suppression of ATPase activity of drDnaA by PprA seems to be an important step in *oriCI* regulation in *D. radiodurans*. Further, drDnaB is found to be a non-specific DNA binding protein with its preference to ssDNA while PprA prefers dsDNA. Therefore, if these differences have accounted to the significant effect of PprA on drDnaA and nearly no effect on drDnaB function cannot be ruled out.

**Fig. 11.**
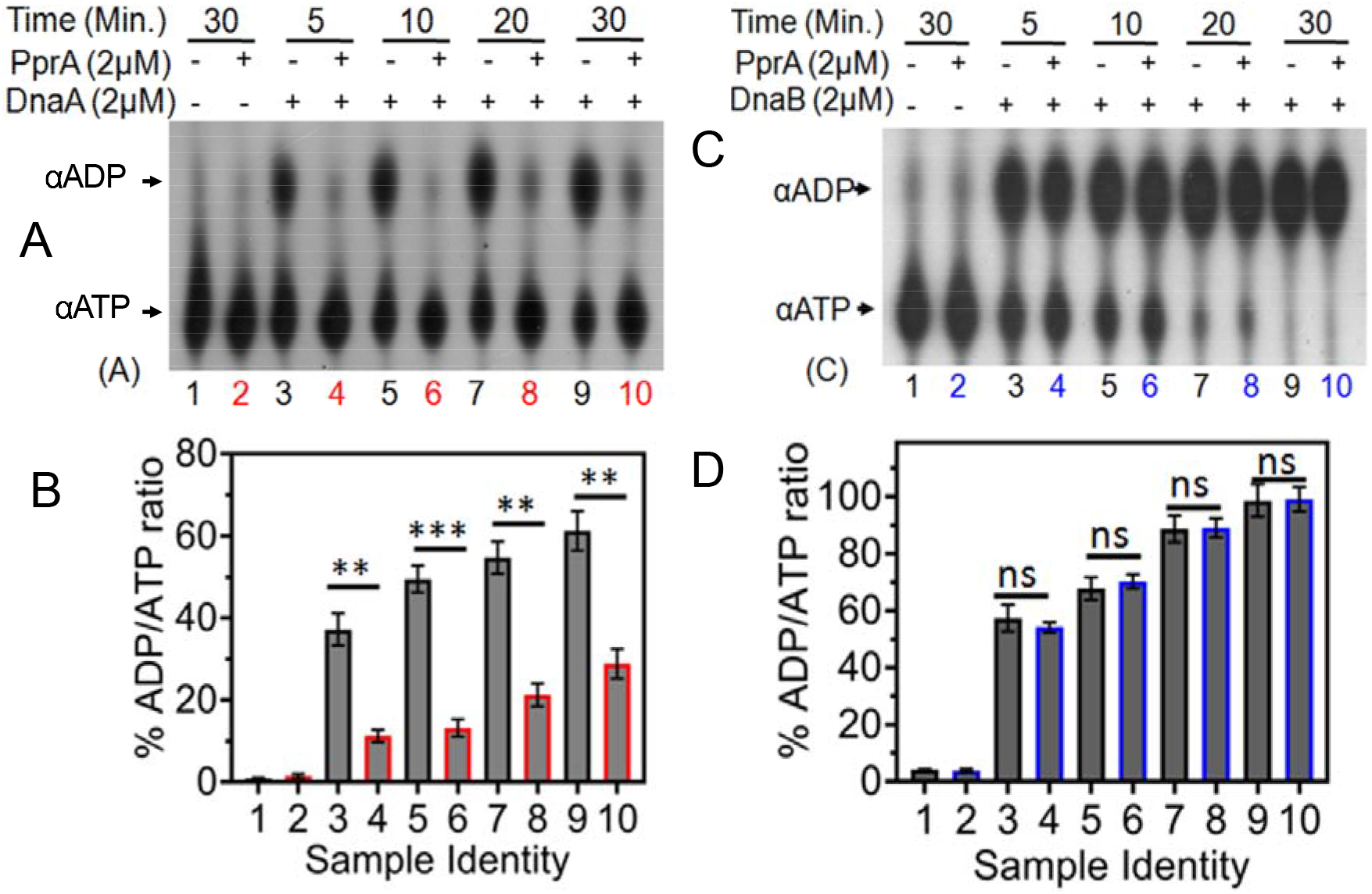
Effect of recombinant PprA on ATPase activity of recombinant drDnaA and drDnaB. Recombinant PprA was incubated with drDnaA (DnaA) (A, B) and drDnaB (DnaB) (C, D) separately. ATP hydrolysis was monitored using [^32^P]-αATP (αATP) at different time interval and generation of [^32^P]-αADP (αADP) product was estimated. Percentage of ADP to ATP ratios were plotted as the mean ± SD (n=3) (B, D). Data sets were analyzed by student t test and significant difference if any is given as the p values less than 0.01 and 0.001 were denoted as (**) and (***), respectively and non-significance is marked as (ns).

**Fig 12.**
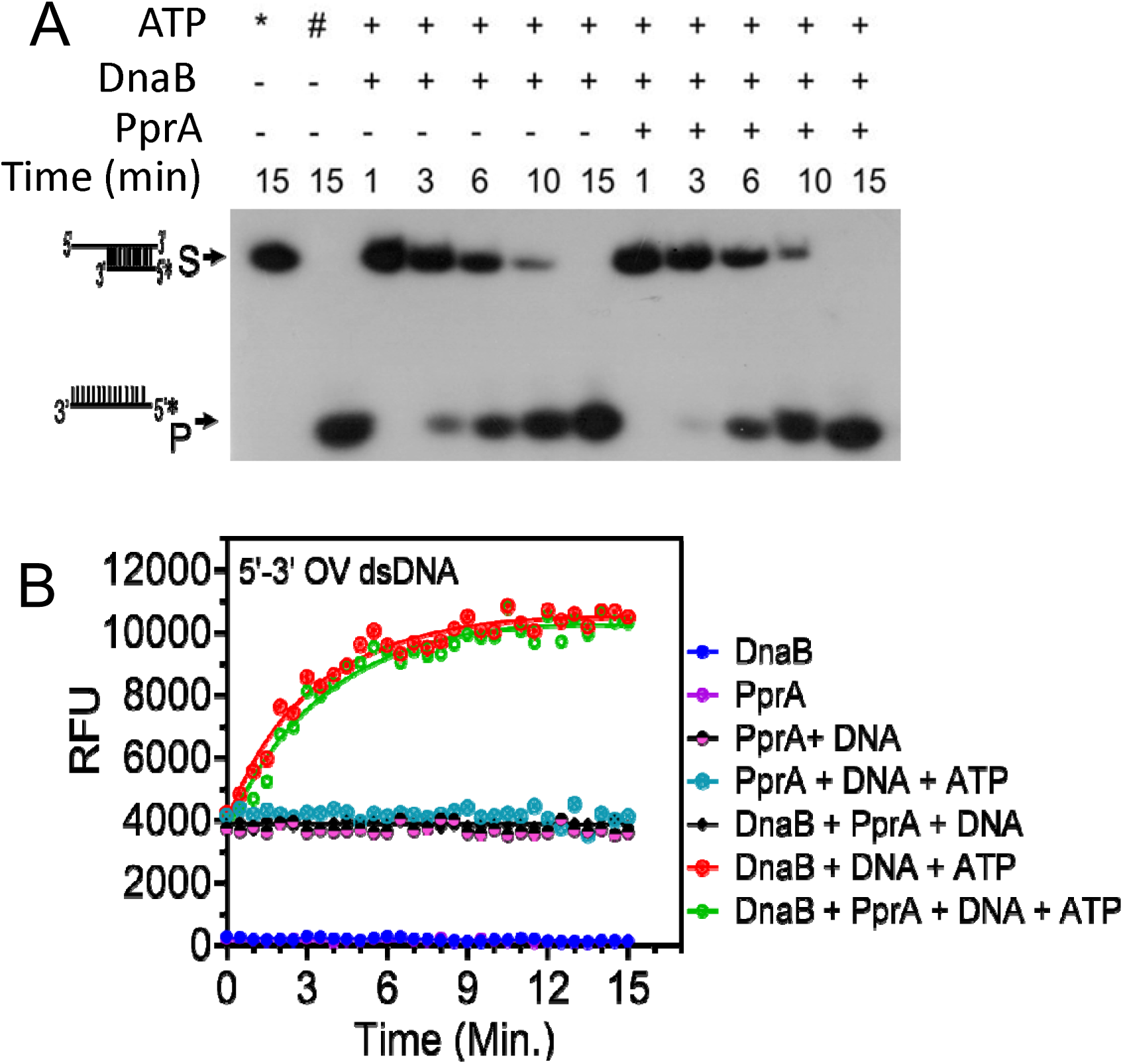
Effect of purified PprA on helicase activity of drDnaB. Recombinant drDnaB (DnaB) was incubated with radiolabeled dsDNA having 5’ overhangs for different time interval in the presence of different combinations of PprA and ATP. The products were analyzed on denaturing PAGE (A). Similarly, FRET substrate of dsDNA with 5’ overhang (Fig 9D) was incubated with drDnaB (DnaB) in different combinations as described in A. Helicase activity was monitored as the loss of FRET and gain of FAM emission with time (B). Data given are representative of the reproducible experiments repeated 2 times.

### PprA negatively regulates drDnaA and drDnaB interaction in surrogate *E. coli*

Since, PprA interact with drDnaA or drDnaB with different affinities, the possibility of PprA affecting drDnaA and drDnaB interactions was hypothesized and tested. The *E. coli* BTH101 cells co-expressing drDnaA and drDnaB interacting combination was co-expressed with PprA from pSpecpprA (Rajpurohit and Misra, 2013) and the expression of β-galactosidase activity was monitored. We observed that the expression of β-galactosidase activity arising from the established interaction of drDnaA and drDnaB (Fig 2) was reduced in the presence of PprA as compared to control without PprA (Fig. 13A-C). This effect of PprA was observed on homotypic and heterotypic interactions of both these replication proteins. These results suggested that PprA interferes with the oligomerization of drDnaA and drDnaB proteins of *D. radiodurans* as depicted in Fig 13D. In different bacteria, it has been reported that oligomerization of both DnaA and DnaB followed by their interaction would be required for faithful DNA replication initiation and elongation. Any defect in oligomerization or interaction of these proteins affects replication initiation process. Therefore, PprA interference in the oligomerization of drDnaA and drDnaB could be a crucial step in the regulation of replication initiation process in *D. radiodurans*.

**Fig 13.**
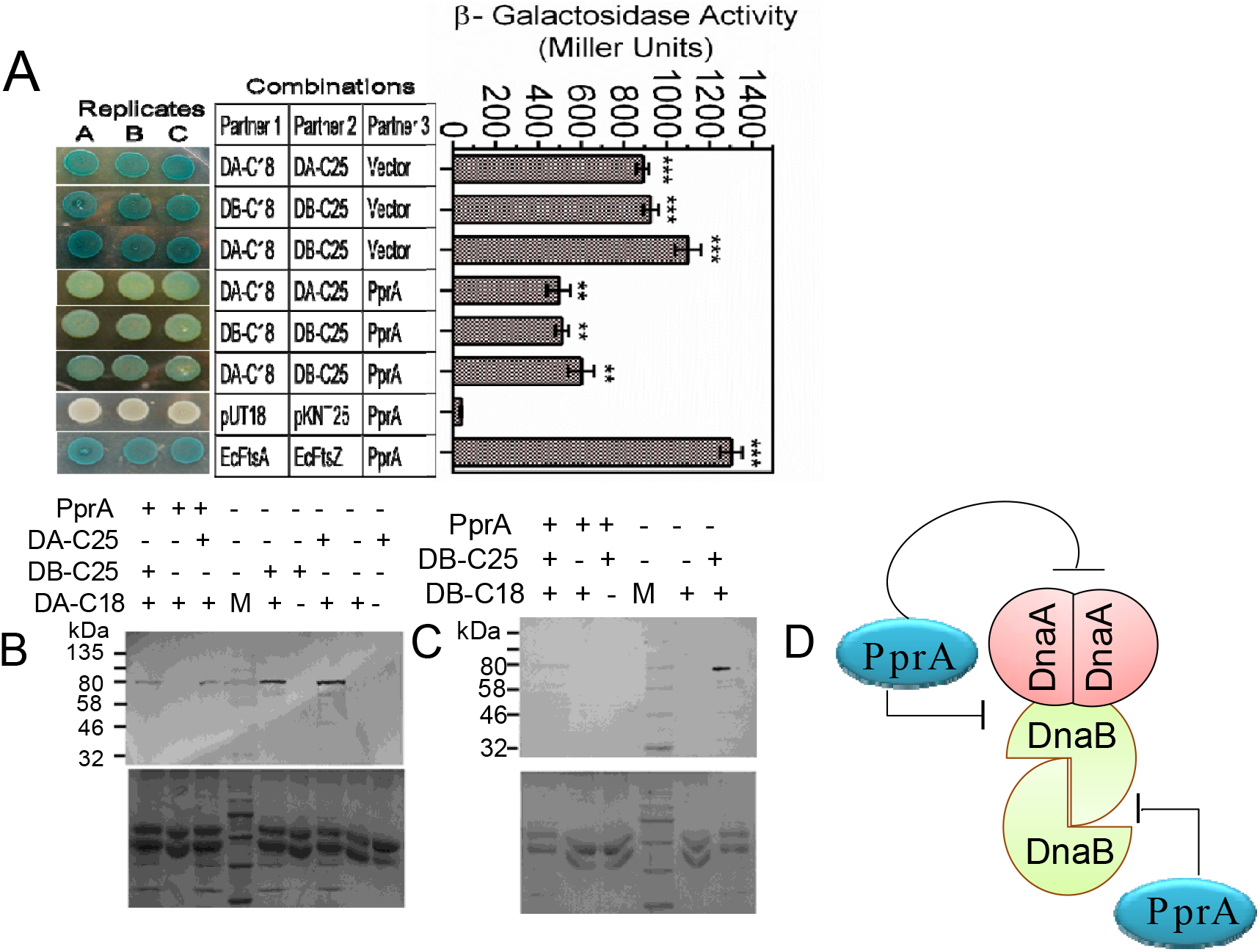
Influence of PprA on drDnaA and drDnaB interaction in surrogate *E.coli*. *E. coli* BTH 101 cells co-expressing C-terminal fusions of T18/T25 tags with drDnaA (DA-C18, DA-C25), DnaB (DB-C25, DB-C18) and PprA in different combinations. The expression of β-galactosidase as an indication of two proteins interaction was monitored in spot test and in liquid assay (A). The pUT18 and pKNT25 vectors were used as negative control while *E. coli* FtsA (EcFtsA) and FtsZ (EcFtsZ) were used a positive control. Total proteins of *E. coli* expressing some of these combinations were immunoprecipitated using T25 antibodies and interacting partners were detected by T18 antibodies (B, C). In Fig. B and C, the upper panels are immunoblots while lower panels are corresponding SDS-PAGE gel. PprA influence on interaction of these proteins is summarised (D).

## Discussion

Molecular basis of *oriCI* regulation has been studied mostly in bacteria that harbour limited copies of single circular chromosome as genetic materials. Very limited information exists on the regulation of DNA replication in bacteria containing multiple chromosomes and ploid genome. In *Vibrio cholerae*, a multipartite genome harbouring bacterium, Chr I replication is initiated by DnaA protein while Chr II replication in initiated by RctB protein that binds and unwinds an array of repeats present in *oriCII* of Chr II (Egan et al., 2005; Jha et al., 2012; Ramachandaran et al., 2017; Bruhn et al., 2018). Since Chr I is larger than Chr II in *V. cholerae*, the temporal regulation of replication initiation could help in synchronization of replication termination of both the chromosomes before the onset of cell division (Rasmussen et al., 2007, Val et al., 2016; Val et al, 2014).

*D. radiodurans*, is another bacterium that harbours polyploid multipartite genome comprising of 2 chromosomes and 2 plasmids (White et al., 1999). Molecular studies on the regulation of DNA replication and control of ploidy of genome elements particularly chromosomes have not been reported yet in this bacterium. Here, for the first time, we have brought forth some evidence to suggest that primary chromosome (chromosome I) encodes the functional DNA replication initiation proteins drDnaA and drDnaB and confers a *cis element* comprised of both *E. coli* type typical DnaA-boxes and non-perfect DnaA-boxes between *drdnaA* and *drdnaN* loci. Based on location on chromosome and structure with 13 AT rich DnaA-boxes, 2 GATC regions and 2 AT rich clusters (12 bp and 20 bp) between 2^nd^ and 3^rd^ DnaA box (White et al., 1999; Luo and Gao, 2018), the cis element was predicted to be *oriC* in chromosome I (*oriCI)* (Fig. 3). Further, we demonstrated that recombinant drDnaA binds specifically to *oriCI* while drDnaB binds preferentially and non-specifically to ssDNA over dsDNA. The presence of ATP has significantly increased the affinity of drDnaA toward *ori CI* by ~ 8 folds (Table 1). Further, the affinity of drDnaA has been reduced with decreasing number of DnaA boxes in *oriCI* (Table 1) and its affinity for non-perfect single DnaA box viz. TTTTCCACA or GTATCCACA was found to be extremely low. The ATPase activity of drDnaA and drDnaB was stimulated by *oriCI* and ssDNA, respectively. Both these proteins undergo homotypic and heterotypic oligomerization *in vitro*. Similar characteristics of DnaA and its interaction with cognate *oriC* have been reported in other bacteria including *B. subtilis* (Yoshikawa & Wake, 1993), *M. tuberculosis* (Zawilak et al., 2004), *T. thermophilus* (Schaper et al., 2000), *Streptomyces coelicolor* and *Spiroplasma citri* (Richter et al., 1998). Like other eubacteria, the C-terminal domain IV of drDnaA is found to be essential for binding with its cognate *oriCI*. The presence of ATPase activity in drDnaA and drDnaB proteins and its stimulation by *oriCI* and ssDNA, respectively, as well as ATP dependent dsDNA helicase function in drDnaB are some of the typical characteristics known for these proteins in other bacteria, and thus make them potentially important for regulation of *oriCI* functions. The DnaBs or its homologs from different bacteria have shown different polarity in dsDNA helicase functions (Caspi et al., 2001; Naqvi et al., 2003). drDnaB shows ATP dependent unwinding of duplex DNA in 5′ → 3′ polarity, which is like the DnaB of *E. coli* (LeBowitz & McMacken, 1986), *H. pylori* (Soni et al., 2003), *M. tuberculosis* (Zhang et al., 2014) and DnaB like helicase in *Bacillus anthracis* (Naqvi et al., 2003). Initiation of DNA replication requires the oligomerisation of DnaA and DnaB at the origin of replication site (Messer, 2002; Zawilak-Pawlik, et al., 2017; Matthew and Simmons, 2019). drDnaA and drDnaB undergo both homotypic and heterotypic oligomerization and interact with *oriCI* region and ssDNA, respectively. These findings together suggest that the 13 AT-rich DnaA-boxes upstream to dr*dna*A is most likely the *origin of replication* in chromosome I, and drDnaA and drDnaB together constitute the replication initiation complex with *oriCI* of *D. radiodurans*.

Various mechanisms have been suggested for the regulation of DNA replication. For instance, in *E. coli* the initiation of DNA replication is regulated by differential affinity of DnaA and SeqA proteins to methylated and hemi-methylated *oriC* and, regulated inactivation of DnaA (RIDA) by Hda (Skarstad and Katayama, 2013). CtrA involvement in the regulation of DNA replication in *Caulobacter crescentus* (Spencer et al., 2009) and *CrtS* protein interaction with methylated DNA in case of *V. cholerae* (Fournes et al., 2018) have been reported. Involvement of genome segregation proteins like Soj and Spo0J in case of *Bacillus subtilis* (Ogura et al., 2003; Murray and Errington, 2008; Scholefield et al., 2011) and ParA and ParB in the regulation of secondary genome copy number in *D. radiodurans* (Maurya et al., 2019a, 2019b) have also been suggested. In case of *oriC* mediated initiation of DNA replication, DnaA binding at *oriC* site triggers the recruitment of (DnaB)6-(DnaC)6 hexameric replicative helicase in several bacteria (Chodavarapu and Kaguni, 2016). Once the helicase is bound to *oriC*, the dissociation of DnaC from DnaB is required for subsequent round of DNA replication. Like many other bacteria, the *D. radiodurans* genome also does not encode DnaC homologue and therefore, the regulation of DNA replication initiation by drDnaA and drDnaB would be either DnaC independent or it will involve some other proteins in the system.

Earlier, it has been shown that ATP hydrolysis by DnaA is required for melting of AT rich region in *oriC* followed by initiation of replication (Messer, 2002; Chodavarapu & Kaguni, 2016). The oligomerisation of both DnaA and DnaB is required for their sequential loading at *oriC*, unwinding of DNA duplex and generation of replication fork, respectively (Arai and Kornberg, 1981; Chodavarapu & Kaguni, 2016; Patel and Picha, 2000). Here, we report that PprA regulates the macromolecular interactions between drDnaA and drDnaB proteins that would be needed for drDnaA to initiate replication. Further, we observed that the genomic content and ploidy levels of *D. radiodurans* cells lacking PprA has increased significantly as compared to wild type (Fig 1). How does PprA regulate genomic content and ploidy levels in *radiodurans* is not clear. However, the higher affinity of PprA to drDnaA (Kd= 5.41 × 10^−7^ ± 1.8 × 10^−8^ [M]) as compared to drDnaB (Kd= 9.71 × 10^−7^ ± 1.1 × 10^−8^ [M]) (Fig 2B, C), inhibition of ATPase activity of drDnaA by PprA while no effect on ATPase and dsDNA helicase function of drDnaB *in vitro* (Fig 11, 12) and the loss of drDnaA and drDnaB interaction in the presence of PprA (Fig 13) are some of the results that indicate the important role of PprA in the regulation of function of replication initiation proteins. Therefore, the absence of PprA if has allowed replication to continue at least one cycle and resulted to an increase in DNA content *per cell* could be speculated. Regulation of DNA replication by a cell cycle regulatory protein competing for binding to *oriC* site has also been suggested (Quoin et al., 1998). Therefore, a possibility of PprA competing with drDnaA binding to *oriCI* was checked and found that PprA does bind to *oriCI* but it was non-specific (Fig S5). These results together supported the role of PprA in regulation of DNA replication perhaps by modulating the activities of replication initiation proteins in *D. radiodurans*.

In conclusion, here we have characterized all the components like *oriCI*, drDnaA and drDnaB involved in *oriC* mediated initiation of bacterial chromosome replication from a multipartite genome harbouring radioresistant bacterium *D. radiodurans.* All these components showed canonical functions required for initiation of chromosomal DNA replication as reported in other bacteria. We found PprA, a protein with pleiotropic functions in this bacterium, regulates the physicochemical properties of these proteins that are required for their functions in *oriC* mediated chromosome replication. While further studies would be required to get the deeper understanding on the role of PprA in regulation of DNA replication in *D. radiodurans* and if it plays a check point regulator of *oriC* mediated DNA replication during DSB repair. The available results that include the increased genomic content per cell in Δ*pprA* mutant and PprA inhibition of drDnaA and drDnaB interaction as well as drDnaA’s ATPase activity together clearly suggest the role of PprA in the initiation of chromosome replication in *D. radiodurans.* How this activity of PprA is regulated under different growth conditions and if phosphorylation of PprA as reported earlier (Rajpurohit et al, 2013) provides a regulatory mechanism in the arrest of DNA replication in quiescent cells engaged in DSB repair during post irradiation recovery of this bacterium would be investigated independently.

## Materials and Methods

### Bacterial strains and plasmids

All the bacterial strains and plasmids and oligonucleotides used in this study are given in Table S1 and Table S2, respectively. *D. radiodurans* R1 (ATCC13939) was grown in TGY (Bactotryptone (1%), Glucose (0.1%) and Yeast extract (0.5%)) medium at 32°C (Schaefer et al 2000). *E. coli* strain NovaBlue was used for cloning and maintenance of all the plasmids while *E. coli* strain BL21(DE3) pLysS was used for the expression of recombinant proteins. *E. coli* strain BTH 101 was used for Bacterial Two-Hybrid System (BACTH) based protein-protein interaction studies. Standard protocols for all recombinant techniques were used as described in (Green and Sambrook, 2012). Molecular biology grade chemicals and enzymes were procured from Merck Inc and New England Biolabs, USA. Radiolabeled nucleotides were purchased from Department of Atomic Energy-Board of Radiation and Isotope Technology (DAE-BRIT), India.

### Cloning, expression and purification of proteins

The dr*dnaA* (DR_0002) and dr*dnaB* (DR_0549) genes were PCR amplified from the genomic DNA of *D. radiodurans* R1 using pETdnaAF & pETdnaAR, and pETdnaBF & pETdnaBR primers, respectively. The PCR products were cloned in pET28a (+) plasmid at *Bam*HI and *Eco*RI sites to yield pETDnaA and pETDnaB plasmids (Table S1). The recombinant plasmids were sequenced, and correctness of inserts was ascertained. The C-terminal (domain IV) truncation of drDnaA was made using pETDAΔCtR primer along with pETdnaAF (Table S2), and resulting plasmid was named pETDAΔCt (Table S1). The recombinant proteins were purified by nickel affinity chromatography as described earlier (Maurya et al., 2019a). Fractions showing more than 95% purity were pooled and dialyzed in buffer A containing 200 mM NaCl and further purified from HiTrap Heparin HP affinity columns (GE Healthcare Life sciences) using a linear gradient of NaCl. Different fractions were analyzed on SDS-PAGE and fractions containing desired protein were pooled and precipitated with 30% w/v ammonium sulphate at 8°C. The precipitate was dissolved in R-buffer (20 mM Tris-HCl pH 7.6, 0.1 mM EDTA, 0.5 mM DTT) containing 1 M NaCl. After centrifugation at 16,000 rpm for 30 minutes, supernatant containing soluble proteins were processed for gel filtration chromatography. The purified protein was dialyzed in dialysis buffer containing 20 mM Tris-HCl pH 7.6, 200 mM NaCl, 50% glycerol, 1 mM MgCl_2,_ 0.5 mM DTT and 1 mM PMSF. Similarly, PprA was expressed from pETpprA and purified as described in (Kota and Misra, 2006) followed by gel filtration as described above.

### Protein DNA interaction studies

The recombinant proteins interaction with different forms of DNA was studied using electrophoretic mobility shift assay (EMSA) as described earlier (Maurya et al., 2019a). In brief, the 500 bp *oriCI* region (1272-1771) containing 13 repeats of 9 mer of DnaA-boxes (T(A/G)TA(T)TCCACA) was PCR amplified using OriIFw and OriIRw primers (Table S1 Fig. 1) and gel purified. In addition, different fragments of *oriCI* with 11, 7, 3 and 1 repeat(s) were also either PCR amplified or chemically synthesised and annealed. DNA substrates were labeled with [γ-^32^P] ATP using T4 polynucleotide kinase. Approximately 30 nM labeled substrate was incubated with different concentrations of recombinant drDnaA in a reaction buffer B containing 50 mM Tris-HCl (pH 8.0), 75mM KCl, 5mM MgSO_4_ and 0.1mM DTT at 37°C for 15 min. The reaction was performed in the presence and absence of 1 mM ATP. For the competition assay, a saturating concentration of protein was incubated with DNA substrate before different concentrations of non-specific competitor DNA of similar length (regions of *ftsZ* gene in Table S1) was added and further incubated as per experimental requirements. Similarly, oligonucleotide 99F (Das and Misra, 2012) was radiolabeled at 5’ end and used as ssDNA substrate for interaction with drDnaB. The interaction of drDnaB with *oriCI* (dsDNA) was monitored in absence and presence of 1 mM ATP as described for drDnaA above. The reaction mixtures were separated on 6-8 % native PAGE gels, the gels were dried, and autoradiograms were developed on X-ray films. The band intensity of unbound and bound fraction was quantified densitometrically and computed using Image J software. The fraction of DNA bound to the protein was plotted against the protein concentrations by using GraphPad Prism6 and the *Kd* values for the curve fitting of individual plots were determined as described in (Maurya et al., 2019a).

### DNA helicase activity assay

The unwinding of double strand DNA substrates (with 5’ overhang as well as 3’ overhang) by drDnaB (2μM) in the presence of different combinations of 1 mM ATP and 2 μM PprA was monitored using fluorescent resonance energy transfer (FRET) approach using 96-well plates as described in (Khairnar et al., 2019). In brief, the substrates used were FAM and BHQ labelled complementary oligonucleotides (Table S1). For preparation of overhang dsDNA, the oligonucleotides of complementary strands producing 5’overhang or 3’ overhang were mixed, heated at 95 ◻C for 10 minutes and then slowly annealed by switching off the heating block for overnight. The helicase assays were performed in volume of 100 μL in helicase buffer (25 mM Tris pH 7.6, 10 mM NaCl, 1 mM MgCl_2,_ 100 μM DTT, 100 μg/ml BSA, 25 mM KCl and 2 % glycerol) at 37 ◻C. To avoid re-annealing of unwound DNA, excess of DNA trap complementary to the BHQ labelled DNA was added to each reaction mixture. The reaction was excited at 490 nm and emission was recorded at 520 nm. The fluorescence signals were monitored at an interval of 30 seconds using microplate reader (Biotek Synergy H1).

Similarly, gel-based DNA helicase activity assay of drDnaB was carried out as descried in (Khairnar et al., 2019). In brief, the 3′ overhang substrate was made by annealing the [^32^P] radiolabelled 3OV99F (5′ TTTTTGCGGTTCATATGGAATTCC3′) with 99F (5′GGAATTCCAATGAACCGCAAAACCGCAAAAACCGTACCGA3′) as described in (Das and Misra, 2012). Similarly, 5′ overhang substrate was generated by annealing of radiolabelled 5OV99F (5′TCGGTACGGTTTTTGCGGTTCATA3’) with 99F oligonucleotides. The annealed substrates were incubated with 2 μM drDnaB with or without 2 μM PprA proteins in the presence and absence of 1 mM ATP or ATP-γ-S in helicase assay buffer at 37°C. Aliquots were taken at different time interval and reactions were stopped with stop solution (20 % glycerol, 20 mM EDTA, 125μg Proteinase K, 0.2 % SDS, 50 mM KCl) at 37 ◻C for 10 minutes. The products were separated on 8 % denaturing PAGE, dried and exposed to X-ray films. The autoradiograms were developed and documented.

### Bacterial Two-Hybrid (BACTH) system assay

drDnaA and drDnaB interaction was monitored in the absence and presence of PprA using bacterial two-hybrid system (BACTH), described in (Maurya et al., 2016, 2018). In brief, the coding sequence of DR_0002 (drDnaA) was cloned at *Bam*HI and *Eco*RI sites in pUT18, pUT18C and pKT25 plasmids to yield pUT18DA, pUT18CDA & pKTDA, respectively. Similarly, coding sequences of DR_0549 (drDnaB) was cloned at *Kpn*I and *Eco*RI sites in pUT18, pUT18C and pKT25 plasmids to yield pUT18DB, pUT18CDB & pKTDB, respectively. The construction of pKNTDA and pKNTDB plasmids has been described in (Maurya et al., 2019b) and pUTpprA in (Kota et al., 2014). *E. coli* strain BTH101 (*cyaA*^−^) was co-transformed with these plasmids in different combinations and expression of β galactosidase was monitored as described earlier. *E.coli* BTH101 harbouring only vectors were used as negative control while those harbouring pUTEFA and pKNTEFZ (Modi and Misra, 2014) were used as positive control. For monitoring the effect of PprA on drDnaA and drDnaB interactions, the PprA was expressed on pSpecpprA (Rajpurohit and Misra, 2013) while p11559 vector was used for negative control. Expression of β-galactosidase as an indication of protein-protein interaction was monitored using spot assay and in liquid culture and β-galactosidase activity was calculated in Miller units as described in (Battesti et al., 2012).

### Co-immunoprecipitation assay

Protein-protein interactions in surrogate *E. coli* (BTH101) and *D. radiodurans* were monitored by co-immunoprecipitation as described in (Maurya et al 2016; Maurya et al., 2018). In brief, the total proteins of the recombinant BTH101 cells co-expressing drDnaA and drDnaB proteins on BACTH plasmids (Table S1) were immunoprecipitated using polyclonal antibodies against T25 and counterpart was detected using T18 antibodies. PprA’s effect on drDnaA and drDnaB interaction was studied by *in trans* expression of PprA on pSpecpprA. Signals were detected using anti-mouse secondary antibodies conjugated with alkaline phosphatase using BCIP/NBT substrates (Roche Biochemical, Mannheim). Similarly, for monitoring drDnaA, drDnaAΔCt and drDnaB in *D. radiodurans* by co-immunoprecipitation, the coding sequences of drDnaA, drDnaAΔCt and drDnaB tagged with polyhistidine were PCR amplified using pETHisFw and pETHisRw primers and cloned in pRADgro plasmid at *Apa*I and *Xba*I sites to yield pRADhisDA, pRADhisDACt and pRADhisDB, respectively (Table S1). In addition, the coding sequences of T18 tagged drDnaA and drDnaB were PCR amplified using BTHF(pv) and BTHR(pv) primers from pUT18DAand pUT18DB plasmids and cloned in p11559 plasmid at *Nde*I and *Xho*I sites to yield pV18DA and pV18DB, respectively (Table S1). These plasmids were co-transformed in different combinations in *D. radiodurans* and induced with 5 mM IPTG as requried. The cell-free extracts of *D. radiodurans* expressing drDnaA, drDnaAΔCt and drDnaB in different combinations were prepared and immunoprecipitated using polyhistidine antibodies as described earlier (Maurya et al., 2019a). The immunoprecipitates were purified using Protein G Immunoprecipitation Kit (Cat. No. IP50, Sigma-Aldrich Inc.) and were separated on SDS-PAGE for western blotting using monoclonal antibodies against T18 as described above. To show interaction of PprA with drDnaA or drDnaB, wild type cells of *D. radiodurans* expressing native PprA were transformed with pV18DA or pV18DB and expression of T18 tagged drDnaA or drDnaB was induced with 5 mM IPTG. The cell free extracts of these transformants were prepared and immunoprecipitated using Anti-PprA antibodies. The purified immunoprecipitates were processed for western blotting using T18 antibodies as described above.

### Surface Plasmon Resonance

Interaction of drDnaA and drDnaB with PprA was also investigated using surface plasmon resonance (SPR; Autolab Esprit, Netherland) as described in (Maurya et al., 2018). For this, 20 μM PprA was immobilized on a bare gold sensor chip using EDC-NHS chemistry as described in user manual at 20°C which results about 200 response units in running buffer [20 mM Tris (pH 7.6)]. The different concentrations (4 - 20 μM) of recombinant drDnaA or drDnaB were used in the mobile phase and incubated with 1 mM ATP and 1 mM MgCl_2_ and passed over the PprA-bound sensor chip in one channel. Reaction buffer containing 20 mM Tris (pH 7.6), 1 mM ATP and 1 mM MgCl_2_ was used as buffer control to flow from another channel over immobilized PprA. The response units for each concentration of proteins were recorded and normalized with buffer control. Further, data were processed using the inbuilt Autolab kinetic evaluation software (V5.4) to find dissociation constant and plotted after curve smoothening using GraphPad Prizm6 software.

### Thin Layer Chromatography for ATPase assay

ATPase activity of drDnaA, drDnaAΔCt and drDnaB was measured as the release of [^32^P] αADP from [^32^P] αATP using Thin Layer Chromatography (TLC) as described earlier (Maurya et al., 2019a). In brief, different concentrations (0 - 3 μM) of drDnaA or drDnaB were mixed with 30 nM [^32^P] αATP in the absence and presence of 0.2 pmol dsDNA or 0.1 pmol ssDNA in 10 μl containing buffer B, respectively. The reaction mixture was incubated at 37^°^C for 30 min. For monitoring the effect of PprA on ATPase activity, 2 μM PprA was used with 2 μM drDnaA or 2 μM drDnaB as described above. The reaction mixture was stopped at different time period (0 - 30 min) with 10 mM EDTA solution and 1 μl of it was spotted on PEI-Cellulose F^+^ TLC sheet. The spots were air-dried, and products were separated on a solid support in a buffer system containing 0.75 M KH_2_PO_4_ / H_3_PO_4_ (pH 3.5). The TLC sheets were air dried, exposed to X-ray film and autoradiograms were developed. Spot intensities of samples were determined by densitometry using Image J 2.0 software. The percentage ratio of ADP to ATP was calculated and plotted using the GraphPad Prizm6 software.

### Fluorescence microscopy

The *D. radiodurans* R1 (WT) and its Δ*pprA* mutant cells were grown till the exponential phase, and the equal number of cells were incubated with 2.5μg 4’,6-diamidino-2-phenylindole dihydrochloride (DAPI) per / ~10^8^ cells for nucleoid staining. These cells were washed twice in phosphate buffer saline and then mounted on slides coated with 0.8% agarose. Cells were imaged in DIC channel (automatic exposure) and DAPI channel (50 milliseconds exposure) under identical conditions for both WT and Δ*pprA* mutant using an Olympus IX83 inverted fluorescence microscope equipped with an Olympus DP80 CCD monochrome camera. Large number of cells (~150) from both WT and Δ*pprA* mutant were processed and fluorescence intensity of DAPI stained nucleoid was measured using ‘intensity profile’ tool in cellSens1.16 software installed with microscope. Mean fluorescence intensity was plotted against sample type using GraphPad Prizm6 software. In addition, relative frequency distribution (%) was also plotted against fluorescence intensity using GraphPad Prizm6 software.

### Determination of ploidy in wild type and Δ*pprA* mutant

The amount of DNA and copy number of all four genome elements like chromosome I, chromosome II, megaplasmid and plasmid per cell of wild type and Δ*pprA* mutant was determined as described earlier (Maurya et al., 2019a). In brief, the exponentially growing wild type and Δ*pprA* cells were adjusted at optical density at 600 nm and their cell numbers were determined using a Neubauer cell counter. The collected cells were washed in PBS followed by 70 % ethanol wash and lysed in solution containing 10 mM Tris pH 7.6, 1 mM EDTA, 4 mg / ml lysozyme. The cells were incubated at 37°C and cell debris was removed by centrifugation at 10000 rpm for 5 min. The integrity of isolated genomic DNA was confirmed by 0.8% agarose gel electrophoresis. DNA content was measured as OD260nm and the genomic copy number was determined using quantitative real-time PCR as described in (Breuert et al., 2006). For quantification of genome copy number, two different genes were taken per replicon showing similar PCR efficiency. For examples, the *ftsZ* and *ftsE* for chromosome I, DR_A0155 and DR_A0002 for chromosome II, DR_B003 and DR_B0076 for megaplasmid and DR_C001 and DR_C018 for small plasmid (Table S2). PCR efficiency of each gene was analysed and was found to be > 96% for each (data not shown). We have followed the Minimum Information for Publication of Quantitative Real-Time PCR Experiments (MIQE) guidelines (Bustin et al., 2009) for qPCR using Roche Light cycler and the cycle threshold (Cp) values were determined. The experiment was performed using three independent biological replicates for each sample. The Cp value for each gene of respective replicon was compared with a dilution series of a PCR product of known concentration, as a standard (Fig S1). The copy number of each replicon by both genes per cell was determined using the cell number present at the time of cell lysis. An average of copy number from two genes per replicon in WT and Δ*pprA* mutant was calculated and represented along with bio-statistical analysis.

## Acknowledgements

Authors are grateful to Prof. V Nagaraja and Prof. D N Rao, Indian Institute of Science, Bangalore, for their intellectual comments on the findings and other valuable suggestions. We thank Ms Shruti Mishra and Dr Sheetal Uppal and Dr C Rajani Kant for their technical comments on the manuscript. GKM and NP are grateful to DAE and DST, Govt of India, for DAE and INSPIRE-research fellowships, respectively.

