## Supplementary material for "PprA interacts with replication proteins and affects their physicochemical properties required for replication initiation in *Deinococcus radiodurans*": Fig S1-S5 and Table S1 -S2

Dr. H. S. Misra

Molecular Biology Division

Bhabha Atomic Research Centre

Mumbai- 400085

$ Present Address: Mahila Mahavidyalaya, Banaras Hindu University, Varanasi-221005. India.

Supplementary figures -	Fig S1- Fig S5

Supplementary Tables - Table S1 and Table S2

**Supplementary figures**

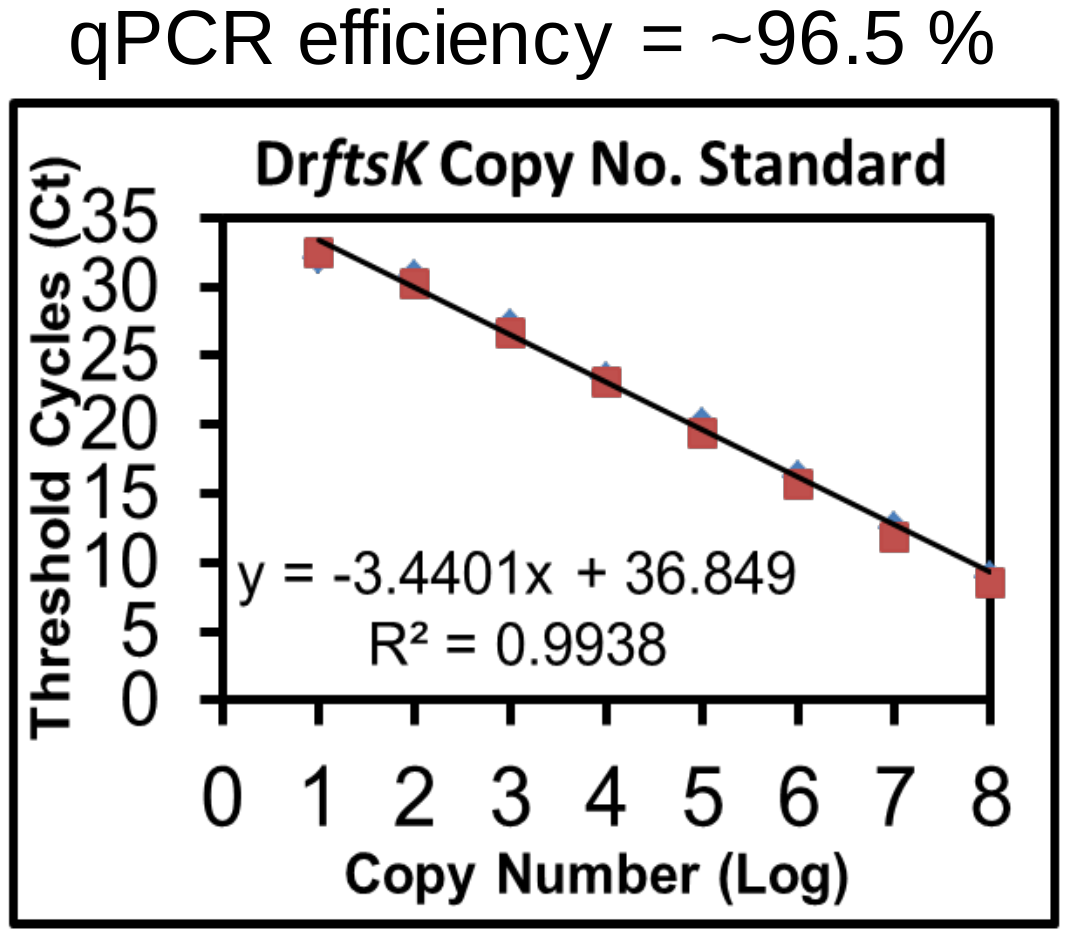

**Fig. S1**. Standard curve for the determination of copy number by RT-PCR. ~300 bp fragment of dr*ftsK* gene was PCR amplified and the known concentrations were taken for qPCR. The obtained Ct value for respective copy number was plotted to give standard curve.

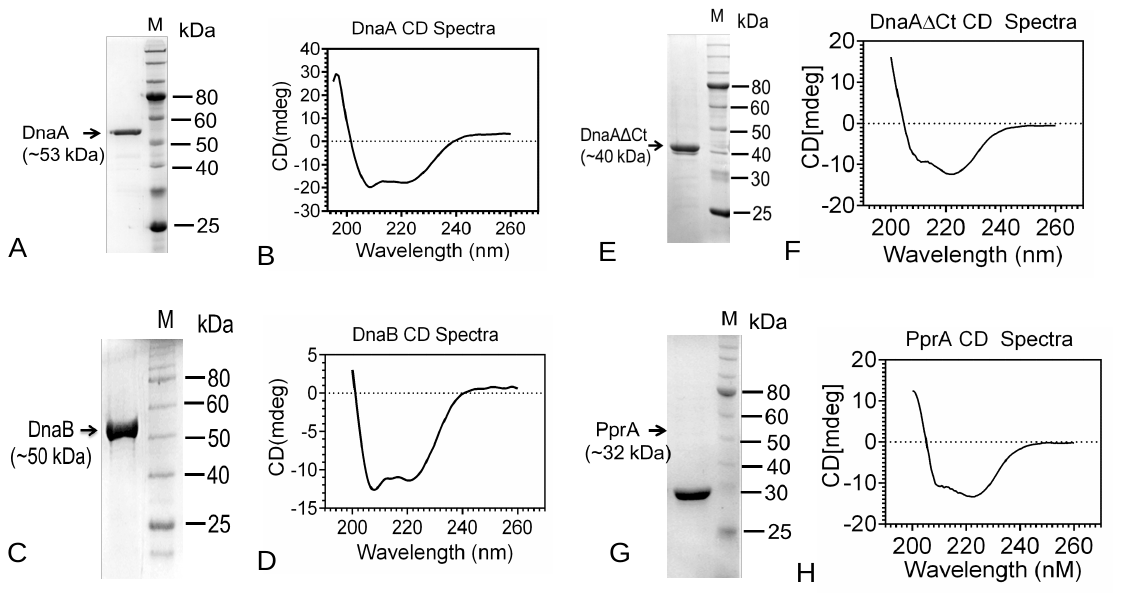

**Fig. S2**. Secondary structure assessment of purified recombinant proteins used in this study. Recombinant drDnaA (A, B), drDnaB (C, D), DnaAΔCt (E, F) and PprA (G,H) were purified from transgenic *E. coli*. These proteins were analysed on SDS-PAGE (A,C,E,G) and by Circular Dichroism analysis (B,D,F,H).

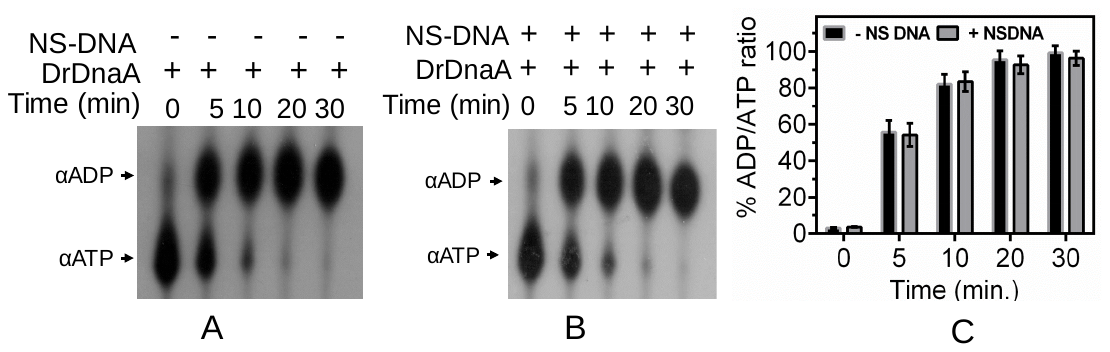

**Fig. S3**. Effect of non-specific DNA on ATPase activity of drDnaA. 2 µM concentration of drDnaA (DnaA) was incubated with [^32^P]-αATP (αATP) at different time interval in the absence (A) and presence (B) of non-specific DNA (NS-DNA) and the generation of [^32^P]-αADP (αADP) product was estimated. Percentage of ADP/ATP ratios were calculated and plotted as mean ± SD (n=3) (C). Results were analyzed using student t-test and significant difference in data set with p values of 0.05 or less is marked with (*) if any.

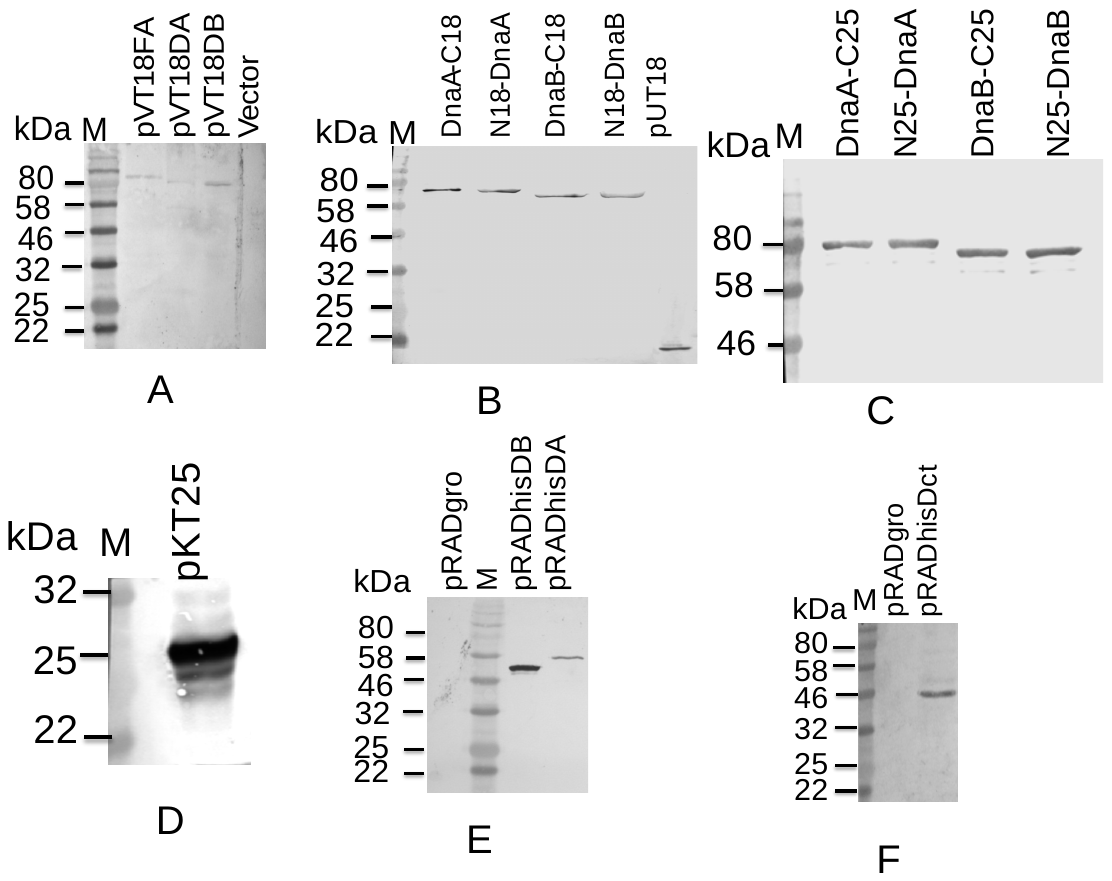

**Fig. S4**. Confirmation of the expression of T18 and T25 fusions of different proteins in *D. radiodurans* (A) surrogate *E. coli* BTH101 (B-D). Expression of polyhistidine tagged fusion of different proteins in *D. radiodurans* (E, F). Fusions were immunoblotted using antibodies against T18 (A,B), T25 (C,D), polyhistidine (E,F).

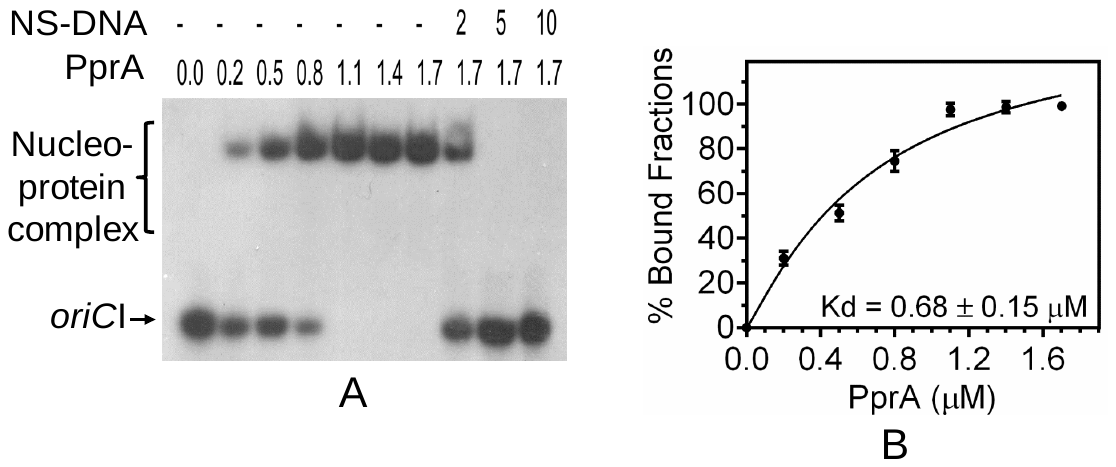
**Fig S5.** PprA binding to *oriCI*. Different concentrations of recombinant PprA (0- 1.7 µM) was incubated with radiolabeled *oriCI*. Saturated reaction mixture at 1.7 µM was chased with 2-fold (2), 5-fold (5) and 10-fold (10) higher molar concentrations of non-specific DNA (NS-DNA). Products were analyzed on non-denaturing PAGE and autoradiograms were developed. Band intensity of free and bound form of DNA substrate was measured densitometrically and mean ± SD (n=3) of percent bound fraction was plotted (B).

**Table S1** – List of bacterial strains and plasmids used in this study

| Bacterial strains | | Genotype | Source | |
| --- | --- | --- | --- | --- |
| *Deinococcus radiodurans* R1 | | Wild type strain ATCC13939 | Lab Stock | |
| *ΔpprA* mutant | | *pprA* gene replaced with chloramphenicol resistance gene cassettes (Cam^R^) | Lab Stock | |
| *E. coli* NovaBlue | | *end*A1 *hsd*R17*(r _K12_ ^−^ m _K12_ ^+^) sup*E44 *thi-1 rec*A1 *gyr*A96 *rel*A1 *lac*F’'*[pro*A^+^B*^+^ lac*I^q^ *Z∆*M15*::*T*n*10 ] (Tet^R^) | NEB Inc., | |
| *E. coli* BTH 101 | | F^-^, *cya-*99*, ara*D139*, gal*E15*, gal*K16*, rps*L1 (Str ^r^)*, hsd*R2*, mcr*A1*, mcr*B1 | Karimova *et al.,* 1998 | |
| *E. coli* BL21(DE3) pLysS | | F–, ompT, hsdS_B_ (r_B_–, m_B_–), dcm, gal, λ(DE3), pLysS, Cm^r^. | Lab stock | |
| **Plasmids** | | | | |
| Sr No. | Plasmids | Characteristics | Sources | MW of  Protein  (~kDa) |
| 1 | pET28a(+) | ~ 5.3 kb plasmid; N-terminal 6XHis tag (Kan^R^) | Novagen | - |
| 2 | pETDnaA | pET28a(+) carrying Dr_0002 at *Bam*HI and *Eco*RI | This study | ~ 53 kDa |
| 3 | pETDnaB | pET28a(+) carrying Dr_0549 at *Bam*HI and *Eco*RI | This study | ~ 50 kDa |
| 4. | pETDAΔCt | pET28a(+) carrying Dr_0002 without domain IV at *Bam*HI and *Eco*RI | This study | ~ 40 kDa |
| 5. | pETpprA | pET28a(+) carrying *pprA* at *Bam*HI and *Hin*dIII | Kota and Misra, 2006 | ~32 kDa |
| 6 | pUT18 | pUC19 derivative, MCS at N-terminal of T18 fragments of adenylate cyclase, ~3 kb, AmpR | Karimova et al., 1998 | ~18 kDa |
| 7 | pUT18C | pUC19 derivative, MCS at C-terminal of T18 fragments of adenylate cyclase, ~3 kb, AmpR | Karimova et al., 1998 | ~18 kDa |
| 8 | pKNT25 | pSU40 derivative, MCS at N-terminal of T25 fragment of adenylate cyclase, ~3.4 kb, KanR | Karimova et al., 1998 | ~25 kDa |
| 9 | pKT25 | pSU40 derivative, MCS at C-terminal of T25 fragment of adenylate cyclase, ~3.4 kb, KanR | Karimova et al., 1998 | ~25 kDa |
| 10 | pUTEFA | pUT18 carrying *E. coli* *ftsA* at *Bam*HI and *Kpn*I | Modi and Mishra, 2014 | ~63 kDa |
| 11 | pKNEFZ | pKNT25 carrying *E. coli* *ftsZ* at *Bam*HI and *Kpn*I | Modi and Mishra, 2014 | ~65 kDa |
| 12 | pRADgro | pRAD1 carrying 261bp *Bgl*II-*Xba*I fragment of promoter (Pgro) from *D. radiodurans* | Misra et al., 2006 | - |
| 13 | pRADhisDnaA | pRADgro carrying N-terminal 6XHis tagged *dnaA* from pETDnaA at *Apa*I*-Xba*I | Maurya et al., 2019b | ~ 53 kDa |
| 14 | pRADhisDACt | pRADgro carrying N-terminal 6XHis tagged dr*dnaA* without domain IV from pETDAΔCt at *Apa*I*-Xba*I | This study | ~40 kDa |
| 15 | pRADhisDB | pRADgro carrying N-terminal 6XHis tagged dr*dnaB* from pETDnaB at *Apa*I*-Xba*I | This study | ~50 kDa |
| 16 | pKNTDA | pKNT25 carrying *drdnaA* at *Bam*HI and *Eco*RI | Maurya et al., 2019b | ~78 kDa |
| 17 | pKTDA | pKT25 carrying *drdnaA* at *Bam*HI and *Eco*RI | This study | ~78 kDa |
| 18 | pUTDA | pUT18 carrying *drdnaA* at *Bam*HI and *Eco*RI | This study | ~71 kDa |
| 19 | pUTCDA | pUT18C carrying *drdnaA* at *Bam*HI and *Eco*RI | This study | ~71 kDa |
| 20 | pKNTDB | pKNT25 carrying *drdnaB* at *Kpn*I and *Eco*RI | Maurya et al., 2019b | ~75 kDa |
| 21 | pKTDB | pKT25 carrying *drdnaB* at *Kpn*I and *Eco*RI | This study | ~75 kDa |
| 22 | pUTDB | pUT18 carrying *drdnaB* at *Kpn*I and *Eco*RI | This study | ~68 kDa |
| 23 | pUTCDB | pUT18C carrying *drdnaB* at *Kpn*I and *Eco*RI | This study | ~68 kDa |
| 24 | pUTpprA | pUT18 carrying *pprA* at *Bam*HI-*Kpn*I | Kota et al., 2014 | ~50 kDa |
| 25 | pVHS559 | An *E. coli* – *D. radiodurans* shuttle plasmid (SpecR) | Lab Stock | - |
| 26 | pSpecpprA | p11559 carrying *pprA* gene at *Nde*I-*Xho*I | Rajpurohit and Misra, 2013 | ~32 kDa |
| 27 | pV18DA | p11559 carrying T18 tagged dr*dnaA* gene from pUTDA at *Nde*I-*Xho*I | This study | ~81 kDa |
| 28 | pV18DB | p11559 carrying T18 tagged dr*dnaB* gene from pUTDB at *Nde*I-*Xho*I | This study | ~78 kDa |

**Table S2-** List of Primers used in this study.

| **Primer** | **Oligonucleotide Sequences** | **Purpose** |
| --- | --- | --- |
| pETdnaBF | 5’GGGGATCCATGGAAACGACTCCGCGT3’ | pETDnaB |
| pETdnaBR | 5’CGGAATTCTCACATCCCCTCCGGCGCCA3’ |  |
| pETDAF | 5’CGGGATCCGTGCGCAAAAACGTCTC 3’ | pETDAΔCt |
| pETDAΔCtR | 5’CGGAATTCTCACATCTCGACCTTGGC3’ |  |
| BTDnaAF | 5’CGGGATCCGGCAGCTGTGCGCAAAAACGTCTC3’ | pUT18DA, pUT18CDA & pKTDA |
| BTDnaAR | 5’GCGAATTCGCGGCCGCCGCCCCGACTTCTTC 3’ |  |
| BTDnaAR+ | 5’GCGAATTCTTACGCCCCGACTTCTTC 3’ |  |
| BTDnaBF | 5’GGGGTACCGGCAGCTATGGAAACGACTCCGCGT3’ | pUT18DB, pUT18CDB & pKTDB |
| BTDnaBR | 5’CGGAATTCGCGGCCGCCATCCCCTCCGGCGCCA3’ |  |
| BTDnaBR+ | 5’CGGAATTCTTACATCCCCTCCGGCGCCA3’ |  |
| BTHF(PV) | 5’GGAATTCCATATGACCATGATTACG 3’ | pV18DA & pV18DB |
| BTHR(PV) | 5’GGCCTCGAGCATATTACTTAGTTA3’ |  |
| pETHisF | 5’AAAAGTACTGGGCCCATGGGCAGCAGCCAT3’ | pRADhisDA, pRADhisDACt, pRADhisDB |
| pETHisR | 5’CGCTTAAGTCTAGATATCTCAGTGGTGGTG3’ |  |
| RTZ Fw | 5'ATCAAGGAATATCTCGA3' | qPCR primers |
| RTZ Rw | 5'CAGCTTTTCGTTGTTCACC3' |  |
| RTEFw | 5’TTGAGCGTCCCCGCGCC3’ |  |
| RTERw | 5’GTGCCCGACGAGGTAGA3’ |  |
| A155RTFw | 5’TTGGCGCATTTTCCCGGC3’ |  |
| A155RTRw | 5’CAGCAGGTTGATCGCCTG3’ |  |
| A0002RTFw | 5’ACCCGGCGTCGTCCAGAG3’ |  |
| A0002RTRw | 5’ GACGACCGGAACTTCAGC 3’ |  |
| B03RTFw | 5’CTGAGTCCTGACGAGTCC3’ |  |
| B03RTRw | 5’TTCCGGGTGACGCAGCAG3’ |  |
| B076RTFw | 5’ATGAGTCCCCCCCTTGCC3’ |  |
| B076RTRw | 5`CGGCGCGATCAACTCCAG3’ |  |
| C01RTFw | 5’ATGTGCTCGCCTCCTAGA3’ |  |
| C01RTRw | 5’ TCACTGTGAAACCTGATC3’ |  |
| C18RTFw | 5’ATGACACAGACGCGGCG3’ |  |
| C18RTRw | 5’GTCCGCGAGGCGCATCAT3’ |  |
| OriIFw | 5’CGGGATCCGTTTTTGCAGCCAACTCCCGA3’ | Primers used for generation of different number of DnaA boxes for EMSA with drDnaA |
| OriIRw | 5’GCGAGCTCCCGTTCCAAATGGCGGTGA3’ |  |
| OriIF1 | 5’CGGGATCCGTAGTAGCAGTAG3’ |  |
| OriIF2 | 5’CGGGATCCCGAAAACCTGATG3’ |  |
| OriI3Fw | 5’AACTTATCCACAGGATATCCACAGGTTTTTCCACAGA3’ |  |
| OriI3Rw | 5‘TTCTGTGGAAAAACCTGTGGATATCCTGTGGATAAGT3’ |  |
| OriI4.1Fw | 5’GGGTTATCCACAGGGC3’ |  |
| OriI4.1Rw | 5’GCCCTGTGGATAACCC3’ |  |
| OriI4.2Fw | 5’GGGTTTTCCACAGGGC3’ |  |
| OriI4.2Rw | 5’GCCCTGTGGAAAACCC3’ |  |
| OriI4.3Fw | 5’GGGGTATCCACAGGGC3’ |  |
| OriI4.3Rw | 5’GCCCTGTGGATACCCC3’ |  |
| Linear 3OVFAM | [FAM]5’GCACTGGCCGTCGTTTTACTCGTGACTGGGAGACCC3’ | FRET substrate  3’ or 5’ overhang Linear dsDNA |
| Linear 5OVFAM | [FAM]5’GCA CTG GCC GTC GTT TTA CTC GTG3’ |  |
| BHQ-comp_ linear OV | 5’TCCAGTCACGAGTAAAACGACGGCCAGTGC3’[BHQ1a~Q] |  |
| TRAP comp linear OV | 5’TACGAGTAAAACGACGGCCAGTGC3’ | FRET TRAP |
| 500bpFw | 5’GGAATTCCATATGCAAGCAGCCAGAATTCGCGT 3’ | PCR amplification of non-specific DNA |
| 500bpRw | 5'CAGCTTTTCGTTGTTCACC3' (RTZRw) |  |
| 260bpFw | 5'ATCAAGGAATATCTCGA3' (RTZFw) |  |
| 260bpRw | 5'CAGCTTTTCGTTGTTCACC3' (RTZRw) |  |
| 170bpFw | 5'ATCAAGGAATATCTCGA3' (RTZFw) |  |
| 170bpRw | 5’CGCATGCGCTTGGGGCCT3’ |  |
| 37bpFw | 5' CGGAATTCGCGGCCGCGATTTTTTCGACACGTTGCAG 3’ (DRparA1R(-SC) |  |
| 37bpRw | 5’CTGCAACGTGTCGAAAAAATCGCGGCCGCGAATTCCG3’ |  |
| 16bpFw | 5’CCGAGGAAAACGTCCA3’ |  |
| 16bpRw | 5’ TGGACGTTTTCCTCGG3’ |  |
